# Astrocytic TDP-43 dysregulation impairs memory by modulating antiviral pathways and interferon-inducible chemokines

**DOI:** 10.1101/2022.08.30.503668

**Authors:** Avital Licht-Murava, Samantha M. Meadows, Fernando Palaguachi, Soomin C. Song, Yaron Bram, Constance Zhou, Stephanie Jackvony, Robert E. Schwartz, Robert C. Froemke, Adam L. Orr, Anna G. Orr

## Abstract

TDP-43 pathology is prevalent in dementia but the cell type-specific effects of TDP-43 are not clear and therapeutic strategies to alleviate TDP-43-linked cognitive decline are lacking. We found that patients with Alzheimer’s disease (AD) or frontotemporal dementia (FTD) have aberrant TDP-43 accumulation in hippocampal astrocytes. In mouse models, induction of widespread or hippocampus-targeted accumulation in astrocytic TDP-43 caused progressive memory loss and localized changes in antiviral gene expression. These changes were cell-autonomous and correlated with impaired astrocytic defense against infectious viruses. Among the changes, astrocytes had elevated levels of interferon-inducible chemokines and neurons had elevated levels of the corresponding chemokine receptor CXCR3 in presynaptic terminals. CXCR3 stimulation altered presynaptic function and promoted neuronal hyperexcitability, akin to the effects of astrocytic TDP-43, and blockade of CXCR3 reduced this activity. Ablation of CXCR3 also prevented TDP-43-linked memory loss. Thus, astrocytic TDP-43 dysfunction contributes to cognitive impairment through aberrant chemokine-mediated astrocytic-neuronal interactions.

**Summary:** In dementia, protein buildup in glia enhances chemokine signaling to synapses and impairs specific aspects of neurocognitive function.

## Introduction

Subcellular mislocalization and dysregulation of transactivating response region (TAR) DNA-binding protein 43 (TDP-43) is a key pathological hallmark of FTD and amyotrophic lateral sclerosis (ALS) (*1-5*). TDP-43 dysregulation is also common in AD and other neurological disorders with pronounced memory loss (*6-10*). Indeed, TDP-43 pathology correlates with cognitive deficits and occurs in up to 50% of AD cases, the majority of hippocampal sclerosis cases, and several other dementias (*11-15*). Despite the prevalence of TDP-43 pathology in various disorders, it is not clear how TDP-43 contributes to disease pathogenesis and cognitive impairments. TDP-43 is a ubiquitously expressed protein that is highly enriched in the nucleus. It is known to regulate RNA processing and transport, among other functions (*1-5, 16-18*). Mislocalization, deficiency, or mutations in TDP-43 can cause pronounced functional deficits and toxicity in animal and cell culture models (*19-22*), indicating that alterations in TDP-43 are sufficient to cause impairments. Recent studies suggest that TDP-43 in glial cells as well as neurons may be dysfunctional and contribute to disease (*7, 11, 23-29*). In mice, selective elimination of mutant TDP-43 from motor neurons delays disease onset, but does not affect disease progression, implicating mutant TDP-43 in non-neuronal cells as a contributor to chronic pathology (*30*). Indeed, astrocyte-targeted expression of TDP-43^M337V^, an ALS-associated mutant form of TDP-43, causes rapid onset of motor deficits and early mortality (*31*). In Drosophila, glia-targeted knockdown of a TDP-43 homologue also causes motor deficits (*32*). Additionally, human induced pluripotent stem cell-derived astrocytes from ALS patients have cell-autonomous TDP-43 accumulation in the cytoplasm (*33*). However, the effects of astrocytic TDP-43 alterations on neurocognitive processes, astrocytic-neuronal interactions, and neuronal activities are not known, and therapeutic targets to alleviate cognitive decline in TDP-43-associated disorders have not been defined.

Here, we investigated whether astrocytic TDP-43 is altered in the human brain and how these alterations influence brain function. We found that TDP-43 accumulates in the cytoplasmic compartment of astrocytes in humans with AD or FTD. In mice, induction of analogous TDP-43 alterations in astrocytes throughout the brain or specifically in the hippocampus caused progressive memory loss that correlated with aberrant changes in antiviral factors expressed by hippocampal astrocytes. In particular, TDP-43 accumulation increased astrocytic levels of interferon-inducible chemokines and presynaptic levels of the corresponding chemokine receptor CXCR3, which promoted neuronal hyperexcitability and memory loss. Thus, dementia-associated TDP-43 alterations in hippocampal astrocytes may contribute to cognitive decline through aberrant engagement of antiviral mechanisms and increases in astrocytic-neuronal chemokine signaling.

## Results

### Astrocytes have extranuclear TDP-43 accumulation in AD and FTD

Neuronal TDP-43 accumulation in the cytoplasm and other alterations have been well-characterized in human samples and model systems. However, TDP-43 pathology is not limited to neurons and might also occur in glial cells, including astrocytes (*34-36*). Astrocytes have crucial roles in brain function and can contribute to disease-related changes, including memory loss, synaptic deficits, and neuroinflammation (*37-39*). Despite robust endogenous expression of TDP-43 in astrocytes, it is not clear if astrocytic TDP-43 is altered in patients with dementia-related memory loss and other neurocognitive impairments. Thus, we first assessed subcellular levels of TDP-43 in astrocytes from AD, FTD, and control (nondementia) cases by performing quantitative immunofluorescent labeling for TDP-43 and the astrocyte marker GFAP in *postmortem* human hippocampal sections. Unlike labeling for other prevalent astrocytic markers, labeling for GFAP is sparser and allows for identification of discrete cells for detailed single-cell neuropathological analyses. Specifically, we measured the levels of TDP-43 immunoreactivity within individual GFAP-positive astrocytic cell bodies and within DAPI-positive nuclei of individual astrocytes. Astrocytes were defined by clear GFAP immunolabeling and characteristic cell morphology. In total, over 800 hippocampal astrocytes were analyzed across groups (**Tables S1**–**S2**). These single-cell subcellular analyses revealed increased levels of diffuse extranuclear TDP-43 immunoreactivity in hippocampal astrocytes from AD cases as compared to age-matched control cases (**Fig. 1A–B**). In contrast, the levels of nuclear TDP-43 immunoreactivity in astrocytes were similar between groups (**Fig. 1C**). Most notably, the ratio of extranuclear-to-nuclear TDP-43 immunoreactivity in astrocytes was increased by approximately 89% in AD cases as compared to controls (**Fig. 1D**). Similar alterations in diffuse astrocytic TDP-43 were also detected in the hippocampus of cases with sporadic or familial FTD (**Fig. 1B–D**), demonstrating that these effects are not unique to AD and are shared among different disorders. It is possible that similar changes in astrocytic TDP-43 accumulation may also occur in other brain regions and other neurological conditions that affect astrocytes.

**Fig. 1.**
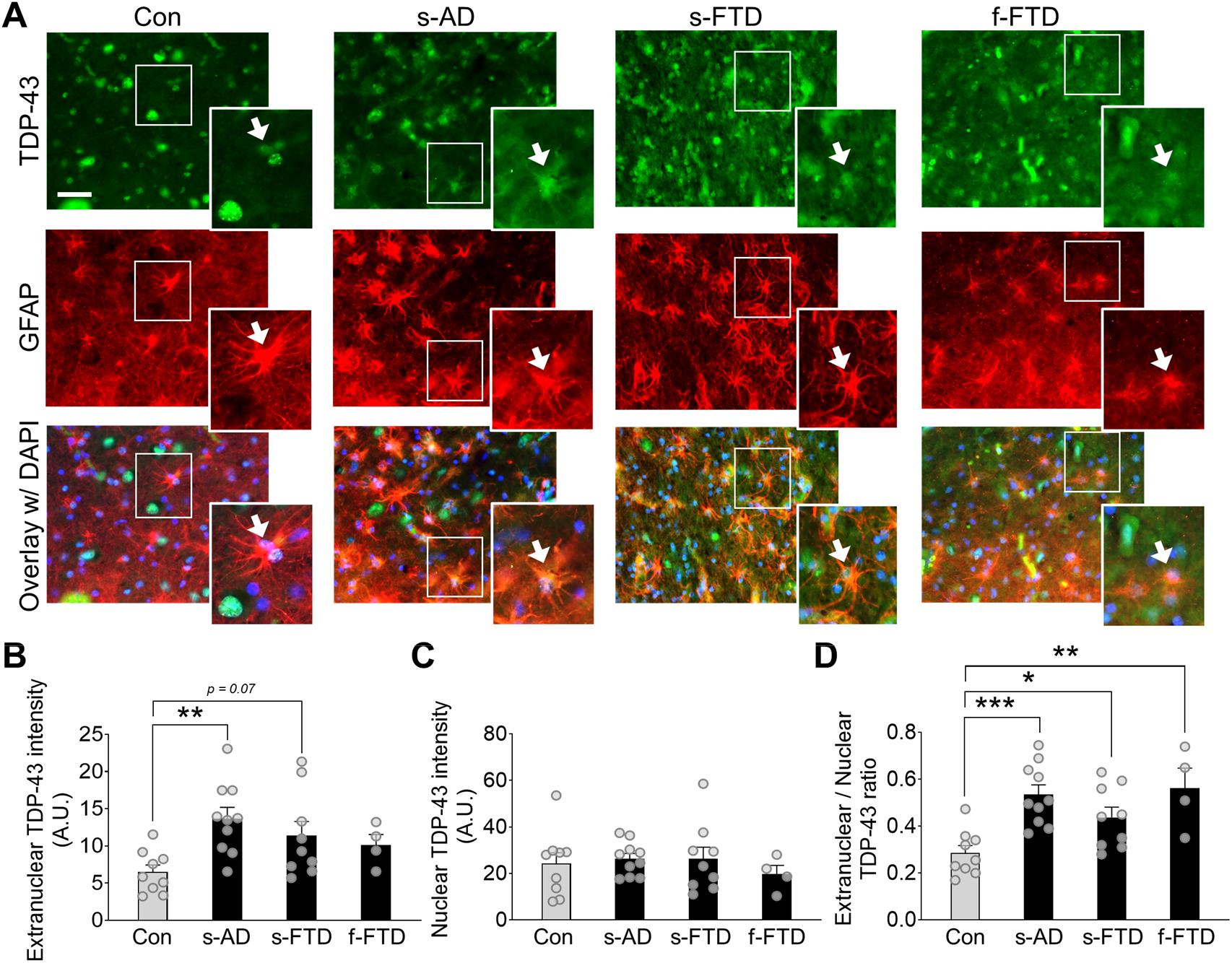
Human astrocytes have increased extranuclear TDP-43 accumulation in AD and FTD. (**A**) Representative images of TDP-43 immunoreactivity (green) in human postmortem hippocampal sections from nondementia controls (Con), sporadic AD (s-AD), sporadic FTD-TDP43 (s-FTD), or familial FTD-TDP43 (f-FTD) cases. The astrocyte marker GFAP (red) was used to visualize astrocytic cell bodies and main processes, and DAPI (blue) was used to visualize cell nuclei within individual astrocytes. Scale bar: 50 μm. (**B**–**D**) Quantification of TDP-43 immunoreactivity within different astrocytic subcellular regions. One-way ANOVA: *F*(3, 28) = 4.21, p = 0.014 (B); *F*(3, 28) = 0.34, p = 0.80 (C); *F*(3, 28) = 7.56, p = 0.0007 (D); Dunnett’s post-hoc test: *p < 0.05, **p < 0.01, ***p < 0.001 vs. Con. n = 9 Con, 10 s-AD, 9 s-FTD, and 4 f-FTD cases.

### Astrocytic TDP-43 alterations cause progressive memory loss

To test whether astrocytic TDP-43 affects behavior and cognitive processes, we generated doubly transgenic mice with inducible and astrocyte-targeted expression of a mutant form of human TDP-43 that accumulates in the cytoplasm (*19*). Specifically, we used the well-validated transgenic *hGFAP*-tTA driver line (*40-44*) to selectively target astrocytes with inducible *tetO*-regulated expression of human TDP-43 that contained a mutated nuclear localization sequence (hTDP43-ΔNLS) (*19*) (**Fig. 2A**). This tet-off system enables suppression of hTDP43-ΔNLS transgene expression using doxycycline (DOX) treatment (*42, 43, 45*). To prevent potential neurodevelopmental effects, breeding pairs and offspring were maintained on a DOX-supplemented diet until weaning at 3 weeks of age, as described previously (*42, 44*). By 3 months of age, widespread astrocytic hTDP43-ΔNLS expression and cytoplasmic accumulation were detected throughout the brain, including the neocortex, hippocampus, thalamus, striatum, and spinal cord in doubly transgenic mice, but not in singly transgenic or nontransgenic controls (**Fig. 2B–C** and **fig. S1**). Expression of hTDP43-ΔNLS was robust in astrocytes, which showed characteristic branching and intermingling with neurons, but was minimal or undetectable in NeuN-positive and Iba1-positive cells (**Fig. 2D–F**).

**Fig. 2.**
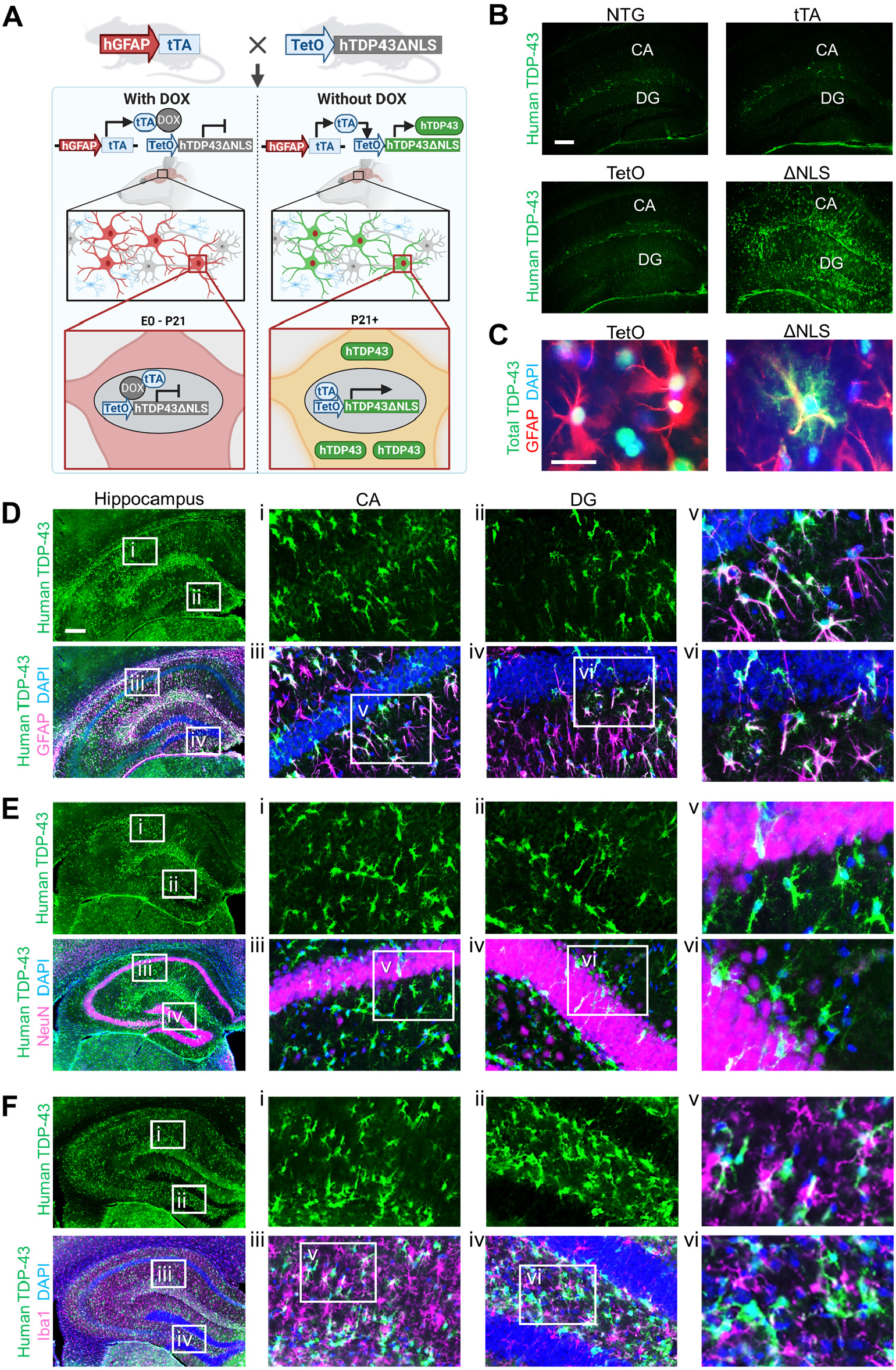
Transgenic mice with inducible TDP-43 alterations in astrocytes. (**A**) Schematic of the tet-off transgenic system used to induce astrocytic expression of human TDP-43 containing a mutated nuclear localization sequence (ΔNLS). (**B**) Representative images of human-specific TDP-43 (green) immunoreactivity in hippocampal sections from 3-month-old nontransgenic (NTG), singly transgenic *hGFAP*-tTA (tTA), and *tetO*-hTDP43-ΔNLS (TetO) control mice, or doubly transgenic ΔNLS mice. Scale bar: 200 μm. (**C**) Representative images of total TDP-43 (green) immunoreactivity in hippocampal sections from TetO controls and ΔNLS mice co-immunolabeled for the astrocyte marker GFAP (red). DAPI (blue) was used to visualize nuclei. Scale bar: 25 μm. (**D**–**F**) Representative images of human-specific TDP-43 (green) immunoreactivity in hippocampal sections from ΔNLS mice co-immunolabeled for the astrocyte marker GFAP (D), neuronal marker NeuN (E), or microglial and macrophage marker Iba1 (F) (magenta). DAPI (blue) was used to visualize nuclei. Insets show magnified views. Scale bars: 300 μm. n = 4–8 mice per genotype.

Previous studies have reported that targeting hTDP43-ΔNLS to neurons causes severe motor deficits and early mortality in mice (*19*) and that an ALS-linked mutant form of hTDP-43 (M337V) expressed in astrocytes is similarly detrimental (*31*). However, we found that targeting hTDP43-ΔNLS to astrocytes did not affect lifespan or alter motor, exploratory, and anxiety-linked behaviors (**fig. S2**). The mice were monitored up to the age of 23–24 months and had normal nesting, burying, and social behaviors, but had mildly increased grooming by 14–15 months of age and increased incidence of ulcerative dermatitis (**fig. S2**). These results highlight the cell type-specific and mutation-specific effects of TDP-43 on brain function and suggest that cytoplasmic TDP-43 accumulation in astrocytes, in contrast to neurons, is not sufficient to cause early mortality, motor deficits, or changes in other innate behaviors.

TDP-43 pathology frequently occurs in patients with AD or hippocampal sclerosis (*6, 11-14, 46*), and TDP-43 pathology correlates with memory impairments (*12, 47, 48*), implicating TDP-43 alterations in memory loss. To determine if astrocytic hTDP43-ΔNLS impairs hippocampus-dependent learning and memory, we tested the mice in the Morris water maze. In this task, mice learn to locate a hidden platform using spatial cues (*49*). At 3–4 months of age, doubly transgenic ΔNLS mice had normal learning during training, and normal memory in a probe test conducted one day after training (**Fig. 3A–B**). However, the mice were moderately impaired in a probe test conducted three days after training (**Fig. 3C**). By 9–10 months of age, doubly transgenic ΔNLS mice had severely impaired performance in probe tests conducted one day and three days after training (**Fig. 3E–H**), but had no deficits in learning or swimming (**Fig. 3D** and **fig. S3**). By 12 months of age, ΔNLS mice also had deficits in novel object recognition, an alternative memory-dependent test, but no changes in total exploration (**Fig. 3I** and **fig. S3**). Thus, astrocytic hTDP43-ΔNLS causes progressive memory deficits but does not markedly impair locomotion or other behavioral functions.

**Fig. 3.**
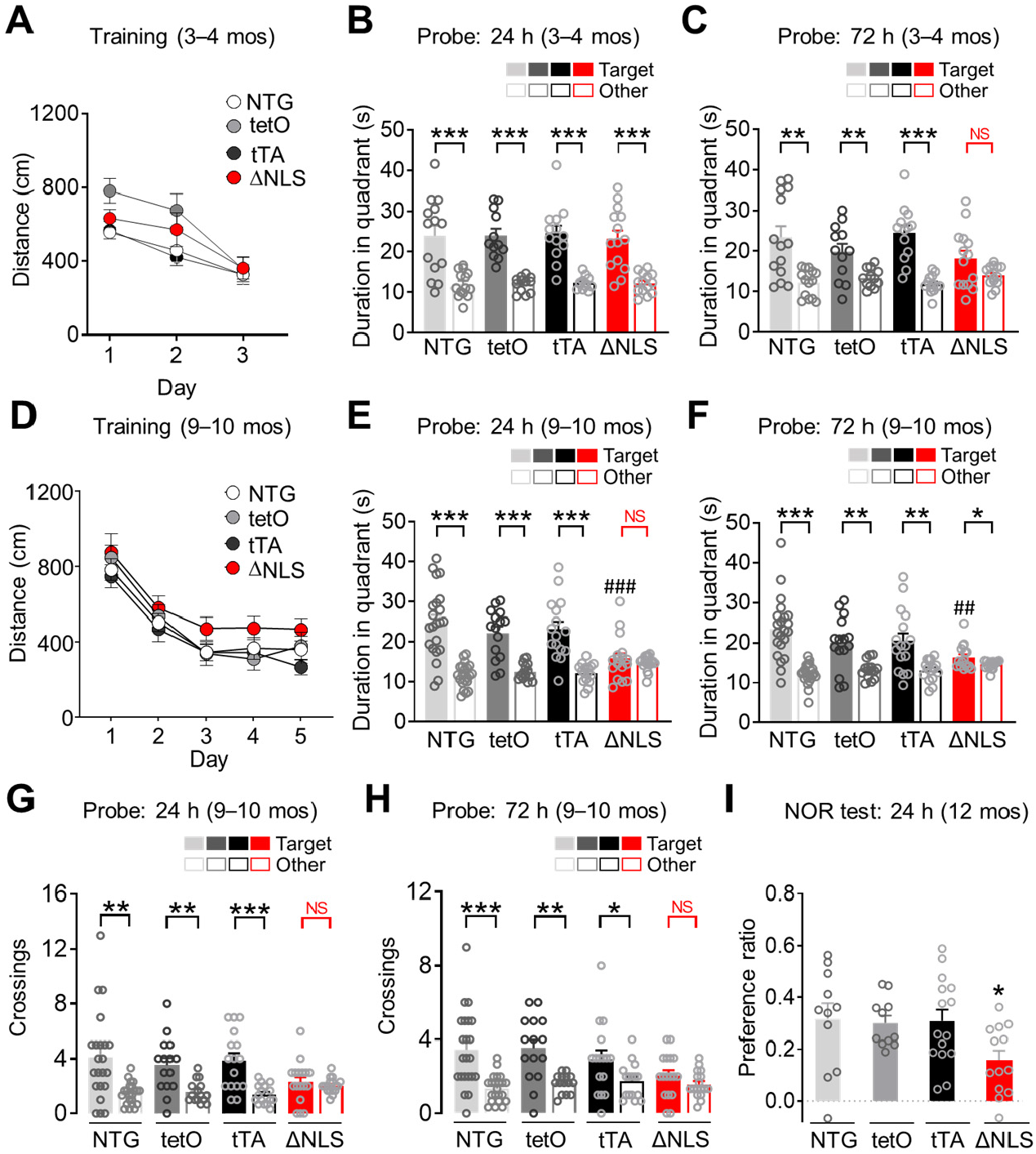
Astrocytic TDP-43 alterations cause progressive memory loss. (**A**–**H**) Nontransgenic controls (NTG), singly transgenic *hGFAP*-tTA (tTA) and *tetO*-hTDP43-ΔNLS (tetO) controls, and doubly transgenic hTDP43-ΔNLS mice (ΔNLS) were tested in the Morris water maze at 3–4 (A–C) and 9–10 (D–H) months of age. (**A**) Distance traveled to reach the platform during hidden platform training (four trials per session, one session per day). Repeated measures two-way ANOVA: *F*(6, 102) = 1.141, p = 0.344 for interaction effect, *F*(3, 51) = 2.635, p = 0.0596 for genotype effect. n = 14 NTG, 12 tetO, 15 tTA, and 14 ΔNLS mice (29 females, 26 males). (**B** and **C**) Probe trials conducted 24 h or 72 h after training. Durations in target and non-target (Other) quadrants. One-way ANOVA (Target): *F*(3, 49) = 0.06677, p = 0.9773 (B); *F*(3, 49) = 1.882, p = 0.1448 (C). Student’s *t* test with Welch’s correction: **p < 0.01, ***p < 0.001 vs. Other. n = 14 NTG, 12 tetO, 15 tTA, and 14 ΔNLS mice (29 females, 26 males). No significant preference for target (NS). (**D**) Distance traveled to reach the platform during hidden platform training (four trials per session, one session per day). Repeated measures two-way ANOVA: *F*(12, 263) = 0.3298, p = 0.9834 for interaction effect, *F*(3, 66) = 2.278, p = 0.0877 for genotype effect. n = 22 NTG, 15 tetO, 16 tTA, and 17 ΔNLS mice (37 females, 33 males). (**E** and **F**) Probe trials conducted 24 h or 72 h after training. Durations in target and non-target (Other) quadrants. One-way ANOVA (Target): *F*(3, 66) = 5.372, p = 0.0023 (E), *F*(3, 65) = 3.548, p = 0.0192 (F); Dunnett’s post-hoc test: ##p < 0.01, ###p < 0.001 vs. NTG Target. Student’s *t* test with Welch’s correction: *p < 0.05, **p < 0.01, ***p < 0.001 vs. Other. n = 22 NTG, 15 tetO, 16 tTA, and 17 ΔNLS mice (37 females, 33 males). (**G** and **H**) Probe trials conducted 24 h or 72 h after training. Crossings of target and non-target (Other) platform locations. Student’s *t* test with Welch’s correction: *p < 0.05, **p < 0.01, ***p < 0.001 vs. Other. n = 22 NTG, 15 tetO, 16 tTA, and 17 ΔNLS mice (37 females, 33 males). (**I**) Novel object recognition test was conducted at 12 months of age. One-way ANOVA and Dunnett’s post-hoc test: *p < 0.05 vs. NTG. n = 14 NTG, 12 tetO, 15 tTA, and 14 ΔNLS mice (29 females, 26 males).

### Astrocytic TDP-43 alterations in the hippocampus are sufficient to cause memory loss

We next examined if the memory deficits induced by astrocytic hTDP43-ΔNLS could be caused by direct effects of astrocytic TDP-43 on the hippocampus, a brain region known to be crucial for memory and susceptible to aging and dementia-associated pathology. To selectively target hippocampal astrocytes *in vivo*, we stereotaxically microinjected adeno-associated viral (AAV) vectors encoding Cre-dependent hTDP43-ΔNLS or hTDP43-WT into the hippocampus of transgenic *Aldh1l1*-Cre mice, which express Cre recombinase predominantly in astrocytes (*50*) (**Fig. 4A**). Although the *Aldh1l1* promoter might be active in other cell types in some contexts and niches of the brain (*51*), recent studies using single-cell transcriptomics and immunolabeling indicate that *Aldh1l1* activity is very low or absent in hippocampal stem cells, including radial glia (*52, 53*). Moreover, *Aldh1l1* promoter-regulated transgene expression is not detected in neuronal progeny (*53*). Nonetheless, to further restrict transgene expression to astrocytes, we also placed the AAV vector-encoded hTDP43-ΔNLS and hTDP43-WT transgenes under the control of the astrocytic promoter *hGfaABC1D* (*54*). This approach created a two-promoter system that requires both *Aldh1l1* and *Gfap* promoter activities for transgene induction. To ensure high efficiency of transgene expression, we used the PHP.eB AAV capsid, which has over 40-fold greater efficiency in transducing neural cells as compared to the standard AAV9 capsid (*55*).

**Fig. 4.**
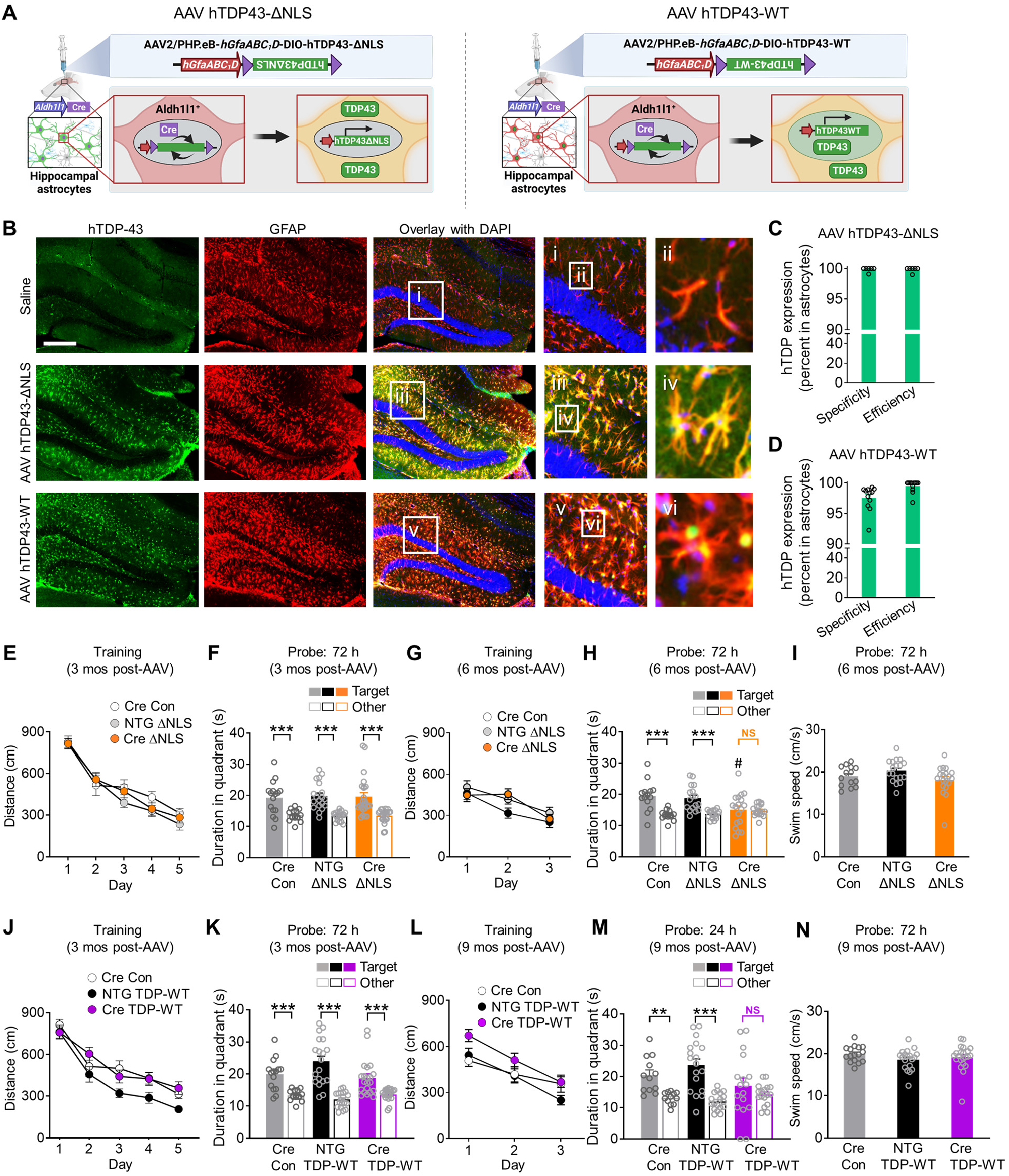
Hippocampus-targeted astrocytic TDP-43 alterations are sufficient to cause progressive memory loss. (**A**) Schematic of hippocampal AAV vector-mediated transgene expression involving a two-promoter system. (**B**) Hippocampal immunolabeling for human-specific TDP-43 (green) and the astrocytic marker GFAP (red). DAPI (blue) was used to visualize nuclei. Yellow indicates overlay of green and red channels. Insets i–vi show magnified views. Mice were assessed for transgene expression 3 weeks after injections. Scale bar: 300 µm. (**C** and **D**) Quantification of hippocampal immunolabeling for human-specific TDP-43. Specificity was defined as percent of hTDP-43-immunoreactive cells in the CA1 and DG layers that were also GFAP-positive; efficiency was defined as percent of GFAP-immunoreactive cells in the CA1 and DG layers that were also hTDP-43-positive. n = 6 AAV hTDP43-ΔNLS and 12 AAV hTDP43-WT-injected *Aldh1l1*-Cre mice. (**E**–**I**) *Aldh1l1*-Cre (Cre) or nontransgenic (NTG) mice were injected at 2–3 months of age and tested in the Morris water maze at 3 or 6 months after injection. n = 16 Cre/Con, 19 NTG/ΔNLS, and 24 Cre/ΔNLS mice (27 females, 32 males) (E–F); n = 14 Cre/Con, 18 NTG/ΔNLS, and 19 Cre/ΔNLS mice (22 females, 29 males) (G–I). (**E** and **G**) Distance traveled to reach the platform during hidden platform training (four trials per session, one session per day). Repeated measures two-way ANOVA: (E) *F*(8, 224) = 0.84, p = 0.57 for interaction effect; (G) *F*(4, 96) = 1.17, p = 0.33 for interaction effect. (**F** and **H**) Probe trials conducted 72 h after training. Durations in target and non-target (Other) quadrants. (H) One-way ANOVA (Target): *F*(2, 46) = 4.18, p = 0.016, Dunnett’s post-hoc test: #p < 0.05 vs. Cre/Con Target. Student’s *t* test with Welch’s correction: ***p < 0.001 vs. Other. No significant preference for target (NS). (**J**– **N**) *Aldh1l1*-Cre (Cre) or nontransgenic (NTG) mice were injected at 2–3 months of age and tested in the Morris water maze at 3 or 9 months after injection. n = 16 Cre/Con, 19 NTG/TDP-WT, and 22 Cre/TDP-WT mice (22 females, 35 males) (J–K), n = 13 Cre/Con, 18 NTG/TDP-WT, and 17 Cre/TDP-WT mice (19 females, 29 males) (L–N). (**J** and **L**) Distance traveled to reach the platform during hidden platform training. Repeated measures two-way ANOVA: (J) *F*(8, 216) = 0.99, p = 0.44 for interaction effect, *F*(2, 54) = 7.19, p = 0.0017 for group effect; (L) *F*(4, 88) = 1.06, p = 0.38 for interaction effect, *F*(2, 44) = 5.06, p = 0.0105 for group effect. (**K** and **M**) Probe trials conducted 24 h or 72 h after training. Durations in target and non-target (Other) quadrants. Student’s *t* test with Welch’s correction: **p < 0.01, ***p < 0.001 vs. Other. (**I** and **N**) Swim speeds during indicated probe trials.

After delivering the vectors intracranially, we performed immunostaining for human TDP-43 protein and markers of astrocytes and other neural cell types. We found that hippocampal transductions with PHP.eB AAV vectors encoding hTDP43-ΔNLS or hTDP43-WT were highly efficient, astrocyte-selective, hippocampus-specific, and stable for months after AAV microinjection (**Fig. 4B–D** and **fig. S4A**–**C**). hTDP43-ΔNLS was localized in astrocyte cell bodies and processes, whereas hTDP43-WT was enriched in astrocytic nuclei. However, similar to hTDP43-ΔNLS, hTDP43-WT was also moderately extranuclear *in vivo* and in isolated astrocytes (**figs. S4B, S4D**–**E**), as described previously in other cell types (*56, 57*), suggesting that both manipulations can cause accumulation of TDP-43 in the cytoplasm.

We next used the Morris water maze to assess learning and memory in AAV-injected transgenic *Aldh1l1*-Cre mice and AAV-injected nontransgenic (NTG) littermate controls. Three months after microinjection, *Aldh1l1*-Cre mice that received AAV encoding hTDP43-ΔNLS had normal learning and probe performance (**Fig. 4E–F**). However, by six months after microinjection, these mice had impaired probe performance as compared to control groups (**Fig. 4G–H**), similar to the results obtained in doubly transgenic hTDP43-ΔNLS mice (**Fig. 3**). NTG mice that received the AAV vector encoding hTDP43-ΔNLS but did not express Cre performed similarly to control *Aldh1l1*-Cre mice that did not receive the AAV vectors, ruling out nonspecific effects by the AAV vectors. Of note, *Aldh1l1*-Cre mice that received the AAV vector encoding hTDP43-WT had normal learning but impaired probe performance by nine months after microinjection (**Fig. 4J–M**), consistent with previous studies showing that accumulation of wild-type TDP-43 is also detrimental (*56-58*). All groups had similar swim speeds (**Fig. 4I** and **4N**).

Additionally, using our two-promoter system, we tested if chronic astrocytic overexpression of other unrelated proteins would similarly cause progressive memory loss. To test this, we transduced hippocampal astrocytes with a PHP.eB AAV vector encoding hM4Di-mCherry (instead of hTDP-43) under the control of *hGfaABC*_*1*_*D* promoter. We did not detect memory deficits in *Aldh1l1*-Cre or NTG mice that received this vector (**fig. S4F**–**K**), further ruling out potential nonspecific effects of AAV transduction and protein overexpression. Together, these results suggest that dysregulation of TDP-43 in hippocampal astrocytes is sufficient to cause progressive memory deficits and that TDP-43 plays essential roles in astrocytic modulation of hippocampal function.

### Astrocytic TDP-43 affects interferon-inducible chemokines and other antiviral factors in a cell-autonomous manner

Astrocytes contribute to memory loss in disease (*42, 59-61*), but the exact mechanisms are not known. In addition, the effects of TDP-43 on astrocytes and astrocytic-neuronal interactions remain unclear. In other cell types, TDP-43 dysfunction can alter inflammatory cascades (*62, 63*), which might promote cognitive decline. Therefore, we next examined if doubly transgenic ΔNLS mice had altered transcription of genes linked to neuroinflammation and glial reactivity in different brain regions that expressed hTDP43-ΔNLS. Targeted transcriptional profiling was carried out in four different brain regions across four genotypes using a microfluidic-based high-throughput RT-qPCR and a custom-designed panel of curated neuroinflammation-related genes. Tau-P301S mice were used as a technical positive control because this model has robust hippocampal gliosis and neuroimmune responses. In contrast to the broad changes in hippocampal gene expression detected in transgenic tau-P301S mice (*64, 65*), we found highly selective changes in gene expression in doubly transgenic ΔNLS mice (**Fig. 5A**). Interferon-inducible chemokines *Cxcl9* and *Cxcl10* were among the top genes affected and were highly increased in the hippocampus, but showed minimal changes in other brain regions, including the neocortex, striatum, and thalamus (**Fig. 5A–C** and **fig. S5A**–**D**). Various markers of astrocytic and microglial reactivity were minimally affected at the RNA and protein levels (**Fig. 5A** and **fig. S5A**–**C, F**).

**Fig. 5.**
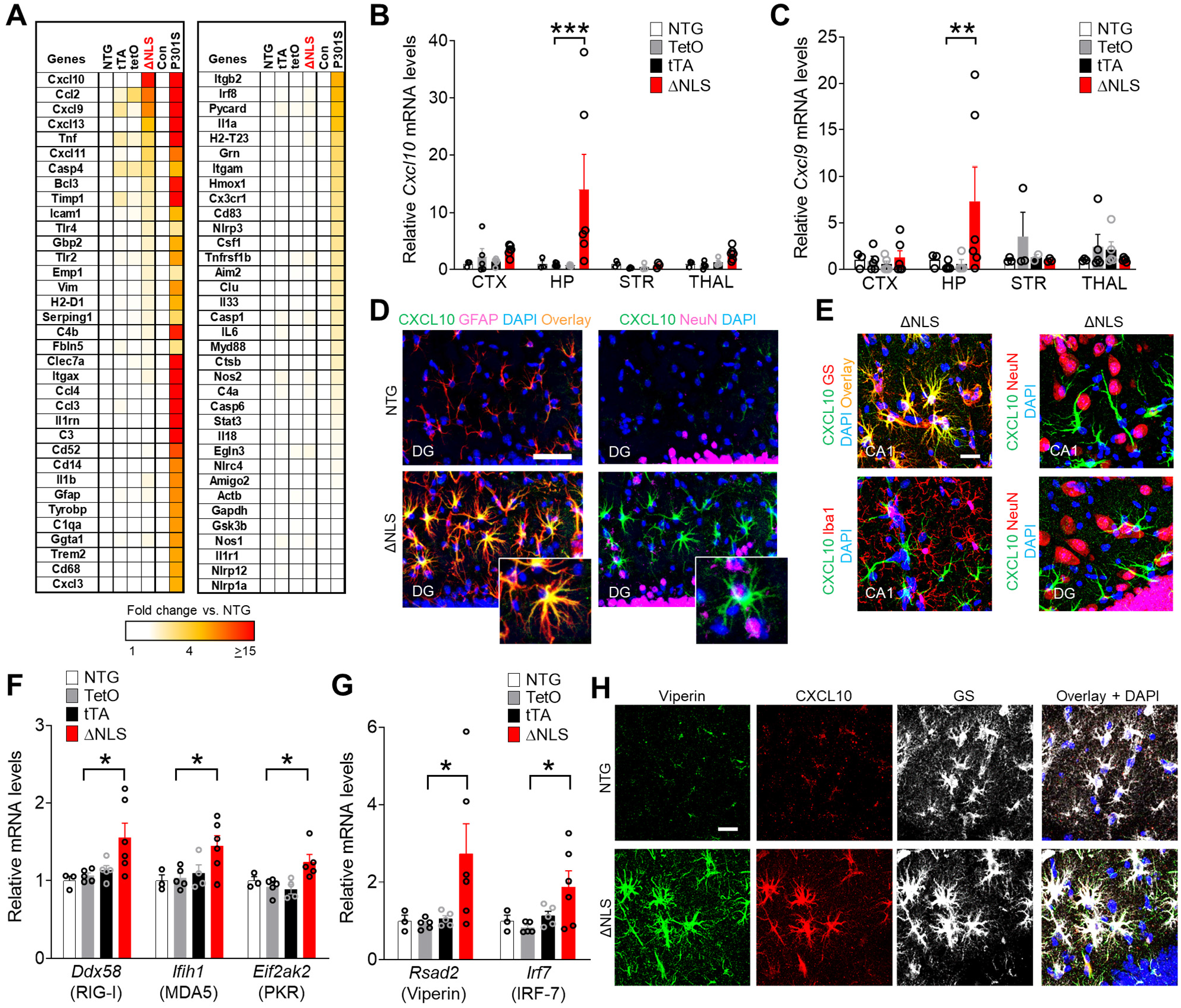
Astrocytic TDP-43 alterations increase hippocampal interferon-inducible chemokines and other antiviral response factors. (**A**) Hippocampal RNA levels for indicated genes in 11-month-old littermate nontransgenic controls (NTG), singly transgenic *hGFAP*-tTA (tTA) and *tetO*-hTDP43-ΔNLS (TetO) controls, and doubly transgenic hTDP43-ΔNLS mice (ΔNLS). Transgenic tau-P301S (P301S) and their littermate controls (Con) at 10 months of age were used for validation and comparison of gene expression. n = 3–6 mice per genotype. (**B** and **C**) RNA levels from indicated brain regions as measured by RT-qPCR. Neocortex (CTX), hippocampus (HP), striatum (STR), and thalamus (THAL). Two-way ANOVA: *F*(9, 52) = 2.06, p = 0.051 for interaction; *F*(3, 52) = 3.061, p = 0.019 for genotype (B); *F*(9, 49) = 1.65, p = 0.13 for interaction; *F*(3, 49) = 0.85, p = 0.47 for genotype (C); Dunnett’s post-hoc test: **p < 0.01, ***p < 0.001 vs. TetO. n = 3–6 mice per genotype and brain region. (**D** and **E**) Representative images of CXCL10 immunoreactivity (green) in the dentate gyrus molecular layer (DG) and CA1 of 11-month-old NTG and ΔNLS mice. Sections were co-immunolabeled for the astrocyte markers GFAP or glutamine synthetase (GS), neuronal marker NeuN, or microglial/macrophage marker Iba1, as indicated. Yellow indicates overlay of green and red channels. DAPI (blue) was used to visualize nuclei. Insets in (D) show magnified views. Scale bars: 100 μm (D), 20 μm (E). (**F** and **G**) Hippocampal RNA levels in 11-month-old NTG controls, singly transgenic TetO and tTA controls, and ΔNLS mice. One-way ANOVA: *F*(3, 15) = 4.19, p = 0.024 (*Ddx58*); *F*(3, 14) = 4.02, p = 0.029 (*Ifih1*); *F*(3, 13) = 5.097, p = 0.015 (*Eif2ak2*); *F*(3, 15) = 3.38, p = 0.046 (*Rsad2*); *F*(3, 15) = 3.29, p = 0.049 (*Irf7*). Dunnett’s post-hoc test: *p < 0.05 vs. TetO. n = 3–6 mice per genotype. (**H**) Representative images of viperin (green), CXCL10 (red), and GS (white) immunoreactivity in the DG of 11-month-old NTG and ΔNLS mice. DAPI (blue) was used to visualize nuclei. Scale bar: 20 μm.

Notably, the increases in CXCL9 and CXCL10 proteins were localized to hippocampal astrocytes, but not neurons, microglia/macrophages, or astrocytes in other examined brain regions (**Fig. 5D–E** and **fig. S5D**–**E**), similar to previous findings of increased astrocytic CXCL10 in a model of multiple sclerosis (*66*) and in humans with AD (*67*). Increases in astrocytic CXCL9 and CXCL10 protein levels were also detected in transgenic *Aldh1l1*-Cre mice that received intrahippocampal injections of PHP.eB AAV vectors encoding hTDP43-ΔNLS or hTDP43-WT (**fig. S5G**), suggesting that astrocytic induction of these chemokines does not require widespread transgene expression and is not unique to the transgenic tet-off system. In addition to CXCL9 and CXCL10, the related chemokine CXCL11 also binds to CXCR3 with high affinity. However, in C57Bl/6 mice (which were used in this study), CXCL11 is not expressed at the protein level due to a premature stop codon (*68, 69*).

Increased expression of CXCL9–11 is typically induced by antiviral or interferon-associated signaling, which regulates glial functions and is linked to cognitive changes, dementia, and neuropsychiatric disorders (*70-77*). One of the primary triggers for antiviral or interferon-associated signaling is activation of pattern recognition receptors (PRRs) that detect non-self or aberrant nucleic acids. Thus, we next tested if sensors of nucleic acids were altered in transgenic mice and isolated astrocytes expressing hTDP43-ΔNLS. Indeed, doubly transgenic mice expressing astrocytic hTDP43-ΔNLS had increased levels of multiple sensors of aberrant dsRNA, including RIG-I (retinoic acid-inducible gene 1; also known as *Ddx58*), MDA5 (melanoma differentiation-associated gene 5; also known as *Ifih1*), and PKR (dsRNA-dependent protein kinase, also known as *Eif2ak2)* (**Fig. 5F**). We also detected increased levels of other antiviral factors, including viperin (*v*irus *i*nhibitory *p*rotein, *e*ndoplasmic *r*eticulum-associated, *in*terferon-inducible; also known as *cig5* or *Rsad2*) and IRF-7 (interferon regulatory factor-7), a master regulator of interferon responses (**Fig. 5G**). Similar to the chemokines, the increases in viperin protein levels were localized to astrocytes but not neurons (**Fig. 5H** and **fig. S5H**). In contrast to the dsRNA sensors, we did not detect changes in the genes encoding dsDNA sensor cyclic GMP-AMP synthase (cGAS) or stimulator of interferon genes (STING) (**fig. S5I**), suggesting that astrocytic TDP-43 alterations may preferentially impact dsRNA sensors and other alternative dsDNA-sensing mechanisms.

Next, we examined whether the changes in astrocytic gene expression were cell-autonomous and linked to functional changes in immune-related signaling. For these experiments, primary astrocytes were isolated from the hippocampus of doubly transgenic ΔNLS or NTG control mice generated in breeding cages with standard chow without DOX. Cell culture purity and transgene efficiency and specificity were evaluated as described in the Methods. As expected, isolated ΔNLS but not NTG astrocytes expressed human TDP43-ΔNLS protein (**Fig. 6A**). Total TDP-43 levels were less than two-fold greater than endogenous TDP-43 levels (**Fig. 6A–B**) and TDP-43 aggregates were not detected (data not shown). Nonetheless, isolated ΔNLS astrocytes had increased levels of multiple interferon-related gene transcripts, including dsRNA-related PRRs (**Fig. 6C**), *Rsad2*/viperin, *Ifnb1, and Ifng* (**Fig. 6D**), suggesting that these gene changes are at least in part cell-autonomous. Isolated ΔNLS astrocytes also had increased levels of phosphorylated NF-κB (**Fig. 6E–F**), supporting the notion that TDP-43 dysfunction affects immune-related pathways in astrocytes, similar to neurons and microglia (*63, 78*).

**Fig. 6.**
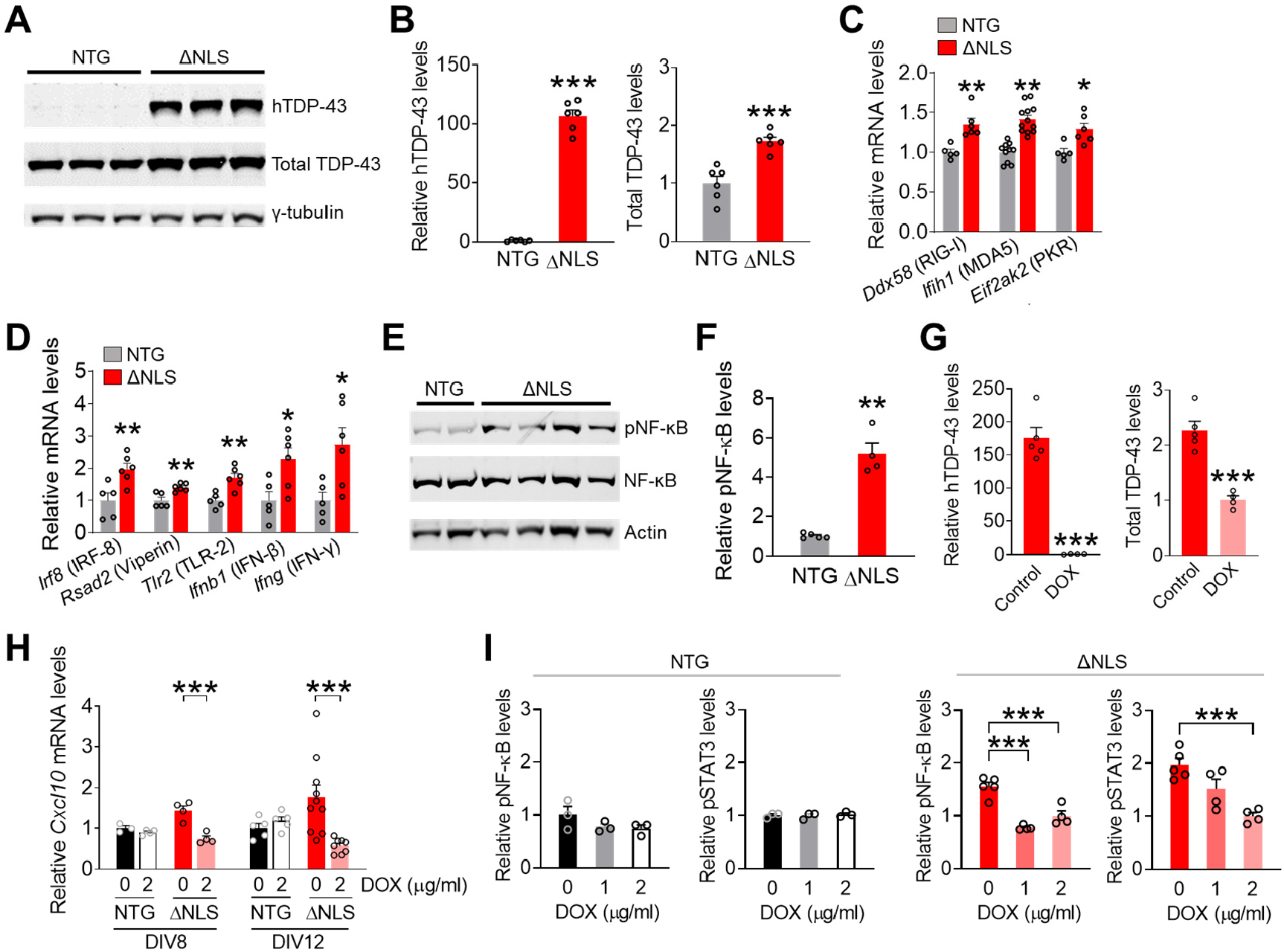
Astrocytic TDP-43 has cell-autonomous effects on antiviral gene expression and neuroimmune pathways. (**A**) Western blots of human TDP-43 or mouse and human TDP-43 levels (total TDP-43) in primary hippocampal astrocytes (DIV 10) derived from nontransgenic (NTG) or doubly transgenic ΔNLS mice. (**B**) Quantification of the Western blots shown in (A). TDP-43 levels were normalized to γ-tubulin levels. Student’s *t* test with Welch’s correction: ***p < 0.001 vs. NTG. n = 6 culture wells per genotype. (**C** and **D**) RNA levels in primary hippocampal astrocytes (DIV 10) were measured by RT-qPCR. Student’s *t* test: *p < 0.05, **p < 0.01 vs. NTG. n = 5–12 wells per genotype. (**E**) Western blots of phosphorylated and total NF-KB levels, and β-actin in primary hippocampal astrocytes (DIV 10) from NTG and ΔNLS mice. (**F**) Quantification of the Western blots shown in (E). Student’s *t* test with Welch’s correction: **p < 0.01 vs. NTG. n = 4–5 wells per genotype. (**G**) Western blot quantification of human-specific TDP-43 and total TDP-43 protein levels in primary hippocampal astrocytes (DIV 10) derived from ΔNLS mice and maintained in DOX (2 µg/ml)-containing media or control media. Student’s *t* test with Welch’s correction: ***p < 0.001 vs. Control. n = 4–5 wells per genotype and treatment condition. (**H**) *Cxcl10* mRNA levels in primary hippocampal astrocytes derived from NTG or ΔNLS mice. Some astrocytes were maintained in DOX-containing media and analyzed at DIV 8 or 12, as indicated. Two-way ANOVA: *F*(1, 11) = 14.62, p = 0.0028 for interaction effect at DIV 8, *F*(1, 24) = 8.28, p = 0.0083 for interaction effect at DIV 12. Bonferroni post-hoc test: ***p < 0.001 vs. 0 DOX per DIV. n = 4–10 wells per genotype and treatment condition. (**I**) Western blot quantification of phosphorylated NF-KB and STAT3 levels in primary hippocampal astrocytes (DIV 10) from NTG or ΔNLS mice and maintained in DOX-containing media or control media, as indicated. Phosphorylated NF-KB and STAT3 levels were normalized to total levels of each protein per sample. One-way ANOVA: *F*(2, 10) = 29.53, p < 0.0001 (pNF-KB in ΔNLS); *F*(2, 10) = 13.68, p = 0.0014 (pSTAT3 in ΔNLS); *F*(2, 6) = 1.77, p = 0.25 (pNF-KB in NTG); *F*(2, 6) = 0.20, p = 0.83 (pSTAT3 in NTG). Dunnett’s post-hoc test: ***p < 0.001 vs. 0 DOX. n = 3–5 wells per genotype and treatment condition.

To further test if these changes were induced by TDP43-ΔNLS expression, some ΔNLS and NTG astrocytes isolated from the hippocampus were maintained in DOX-supplemented media. As expected, DOX-treated ΔNLS astrocytes had minimal expression of human TDP-43 as compared to ΔNLS astrocytes maintained without DOX (**Fig. 6G**). DOX-treated ΔNLS astrocytes had reduced levels of *Cxcl10* gene expression (**Fig. 6H**) and reduced levels of phosphorylatedNF-KB and STAT3 (**Fig. 6I**) as compared to astrocytes maintained without DOX. These signaling factors were not affected in NTG control astrocytes treated with DOX. Thus, TDP-43 altered baseline astrocytic immune-related gene expression and signaling.

Although the approximately two-fold increases in total levels of TDP-43 induced moderate changes in gene expression and signaling, these changes were chronic and associated with marked functional effects on antiviral responses to pathogens. To examine these effects, we used a mimic of viral dsRNA, polyinosinic-polycytidylic acid (poly(I:C)), as a positive control for induction of antiviral responses. Transfection with poly(I:C) induces robust astrocytic antiviral responses at least in part via TLR3 and MDA5 (*79*). Indeed, isolated astrocytes acutely transfected with poly(I:C) had robust induction of antiviral genes and reduced levels of infection by vesicular stomatitis virus (VSV), a negative-sense RNA virus (**Fig. 7A** and **fig. S6A**). In comparison to NTG controls, ΔNLS astrocytes had increased levels of VSV infection (**Fig. 7A**), suggesting that antiviral responses were impaired. ΔNLS astrocytes also had increased infection by adenovirus (**Fig. 7B**), a double-stranded DNA virus. Cell culture density was similar between treatments and genotypes (**fig. S6B**–**C**), indicating that the observed increases in viral infections were unlikely to result from altered cell survival or density. We also tested if astrocytes were more susceptible to herpes simplex virus-1 (HSV-1), a highly neurotropic double-stranded DNA virus increasingly implicated in AD (*80-82*). Indeed, ΔNLS astrocytes had increased levels of HSV-1 infection (**Fig. 7C–E**). Thus, in addition to affecting cognitive function, alterations in astrocytic TDP-43 impaired antiviral defenses, which may predispose the cells to infectious pathogens. However, further studies are necessary to address how astrocytic proteinopathy affects host responses to microbial pathogens and possibly other neuroimmune challenges, and whether alterations in type I and/or type II IFN pathways and other mechanisms are contributing factors.

**Fig. 7.**
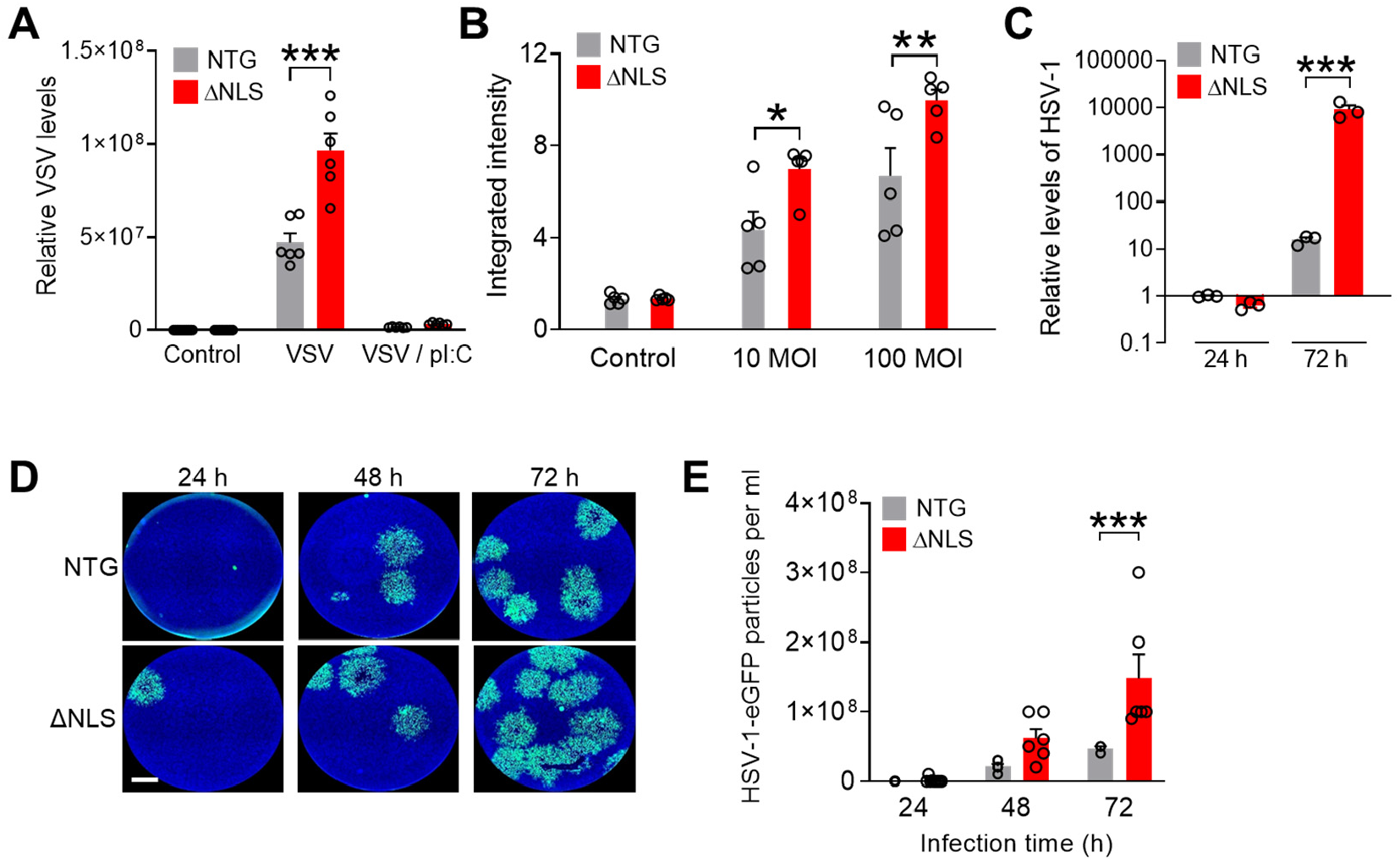
Astrocytic TDP-43 affects antiviral defenses in a cell-autonomous manner. (**A**) Primary astrocytes (DIV 9) from NTG and ΔNLS mice were infected with vesicular stomatitis virus (VSV,100 MOI) for 24 h. Some wells were also transfected with poly(I:C) (pI:C). VSV levels were measured by RT-qPCR. Two-way ANOVA: *F*(2, 41) = 34.64, p < 0.0001 for interaction effect; *F*(1, 41) = 40.60, p < 0.0001 for genotype effect. Bonferroni post-hoc test: ***p < 0.001 vs. NTG. n = 6–12 wells per genotype and treatment condition. (**B**) Primary astrocytes (DIV 9) from NTG and ΔNLS mice were infected with adenovirus tagged with eGFP at indicated MOIs for 24 h. eGFP levels were measured by quantitative microscopy. *F*(2, 24) = 3.47, p = 0.047 for interaction effect; *F*(1, 24) = 13.73, p = 0.0011 for genotype effect. Bonferroni post-hoc test: *p < 0.05, **p < 0.001 vs. NTG. n = 5 wells per genotype and treatment condition. (**C**) Primary astrocytes (DIV 8) from NTG and ΔNLS mice were infected with HSV-1 tagged with eGFP (0.01 MOI) for 24 h or 72 h. gB DNA levels were normalized to 18S DNA per sample to DNA. Two-way ANOVA: F(1, 12) = 15.66, p = 0.0019 for genotype effect; F(2, 12) = 9.225, p = 0.0037 for interaction effect. Bonferroni’s post-hoc test: ***p = 0.0003 vs. NTG-72 h. n = 3 wells per genotype and treatment condition. (**D** and **E**) Conditioned media was collected from primary NTG or ΔNLS astrocytes after infection with HSV-1-eGFP (0.01 MOI). Astrocytes were washed 3 h after infection and conditioned media was analyzed after indicated durations using the plaque assay in Vero cells. (**D**) Representative images of Vero cells after treatment with conditioned media from NTG or ΔNLS astrocytes that were infected with HSV-1-eGFP for indicated durations. Scale bar: 1200 µm. (**E**) Number of viral particles in conditioned media from HSV-1-infected NTG or ΔNLS astrocytes. Two-way ANOVA: F(1, 33) = 15.24, p = 0.0004 for genotype effect; F(2, 33) = 5.58, p = 0.0082 for interaction effect. Bonferroni’s post-hoc test: ***p = 0.0009 vs. NTG-72 h. n = 3–9 wells per genotype and treatment condition.

### Astrocytic TDP-43 alterations are linked to increased presynaptic levels of the chemokine receptor CXCR3 and CXCR3-mediated neuronal impairments

Given that memory was impaired in ΔNLS mice, we next explored how changes in astrocytic interferon-inducible factors might affect neuronal functions. Upon release, interferon-inducible chemokines CXCL9–11 activate the shared G protein-coupled receptor CXCR3 (*83*), which can be expressed by neurons (*67, 84*), microglia (*85*), and potentially other cell types. Notably, patients with FTD or AD show increased levels of CXCL10 (*86*) and have hippocampal CXCR3 expression predominantly in neurons (*67*), suggesting that neuronal CXCR3 might play a role in these disorders. We found that hippocampal *Cxcr3* RNA and protein levels were increased in ΔNLS mice (**Fig. 8A–C**) and the majority of CXCR3-immunoreactive puncta were localized in neurons (**Fig. 8D–E**). However, ΔNLS mice did not have increased levels of CXCR3 immunoreactivity within neuronal cell bodies (**Fig. 8D–E**). Co-immunolabeling for CXCR3 and different presynaptic and postsynaptic markers revealed that ΔNLS mice had two-fold increases in CXCR3 selectively within synaptophysin-positive puncta, but not within PSD-95 or gephyrin-positive puncta (**Fig. 8F–J**), suggesting that the increases in hippocampal CXCR3 levels were localized to presynaptic terminals.

**Fig. 8.**
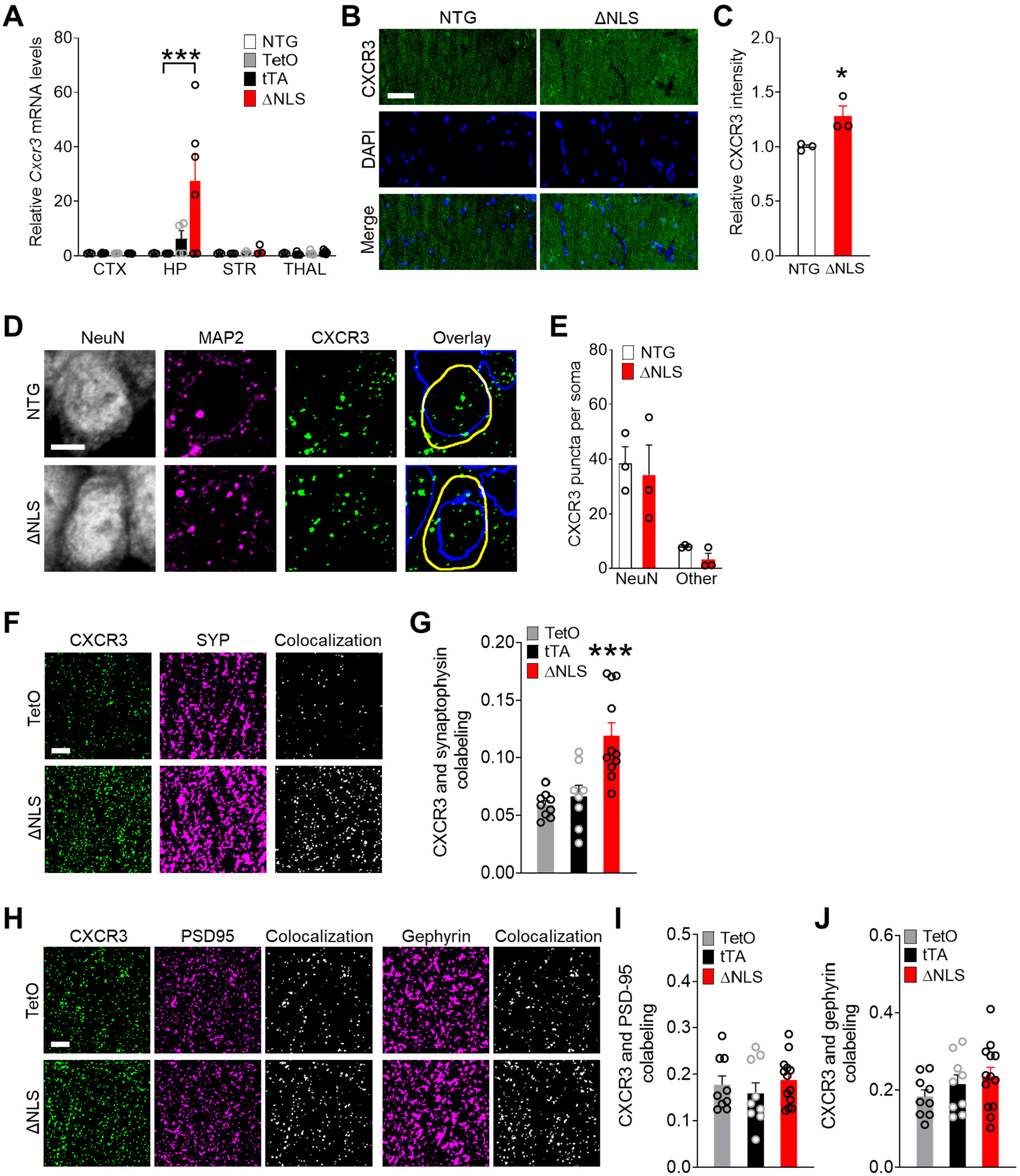
Astrocytic TDP-43 increases chemokine receptor CXCR3 levels in presynaptic terminals. (**A**) *Cxcr3* RNA levels from indicated brain regions of NTG, singly transgenic controls (TetO and tTA), and doubly transgenic ΔNLS mice. Neocortex (CTX), hippocampus (HP), striatum (STR), and thalamus (THAL). Two-way ANOVA: *F*(9, 48) = 2.89, p = 0.0081 for interaction; *F*(3, 48) = 2.93, p = 0.043 for genotype; Dunnett’s post-hoc test: ***p < 0.001 vs. TetO. n = 3–5 mice per genotype and brain region. Representative images (**B** and **D**) and quantification (**C** and **E**) of hippocampal immunoreactivity for CXCR3 in the CA1 radiatum parenchyma (B–C) or specifically in CA1 neuronal cell bodies (D–E) as delineated by co-immunolabeling with neuronal marker NeuN versus non-NeuN regions in NTG and ΔNLS mice. Neuronal nuclei are indicated by blue traces; cell somas are indicated by yellow traces. Arbitrary fluorescence intensity units were normalized to NTG mice (C). Student’s *t* test: *p = 0.039; n = 3 mice per genotype. (**F** and **G**) Colocalization of CXCR3 and the synaptic marker synaptophysin in the CA1 region of singly transgenic controls and doubly transgenic ΔNLS mice. Mander’s overlap coefficient was used to assess colocalization. One-way ANOVA: *F*(2, 25) = 12.94, p < 0.0001; Dunnett’s post-hoc test: ***p < 0.001 vs. TetO. n = 8–11 mice per genotype. (**H**–**J**) Colocalization of immunoreactivity for CXCR3 and the synaptic markers PSD-95 (H, I) or gephyrin (H, J) in the CA1 region of singly transgenic controls and doubly transgenic ΔNLS mice. Mander’s overlap coefficient was used to assess colocalization. n = 9–13 mice per genotype. Scale bars: 50 µm (B), 5 µm (D, F, H).

CXCR3 activation triggers G_i/o_-coupled signaling, which can inhibit presynaptic neurotransmitter release (*87, 88*). However, the presynaptic effects of CXCR3 have not been previously defined. Thus, we next assessed whether activation of presynaptic CXCR3 affects neuronal activity by using the multi-electrode array system. For these experiments, primary mouse neurons were transduced with PHP.eB AAV Syn*-Cx*cr3-2HA-neurexin1α. In this vector, the neurexin-1α sequence targets CXCR3 to presynaptic terminals, which simulates the presynaptic enrichment of CXCR3 observed in ΔNLS mice and limits artificial effects of CXCR3 on neuronal excitability (*89*). We confirmed that the vector was functional in neurons and that CXCR3 stimulation with chemokines induced characteristic intracellular signaling (**fig. S7A**–**B**). We found that acute chemokine treatment inhibited spontaneous neuronal activity within minutes of treatment (**Fig. 9A**), suggesting that CXCR3 rapidly suppresses neuronal firing, possibly through modulation of calcium channels and presynaptic vesicles (*87, 88*).

**Fig. 9.**
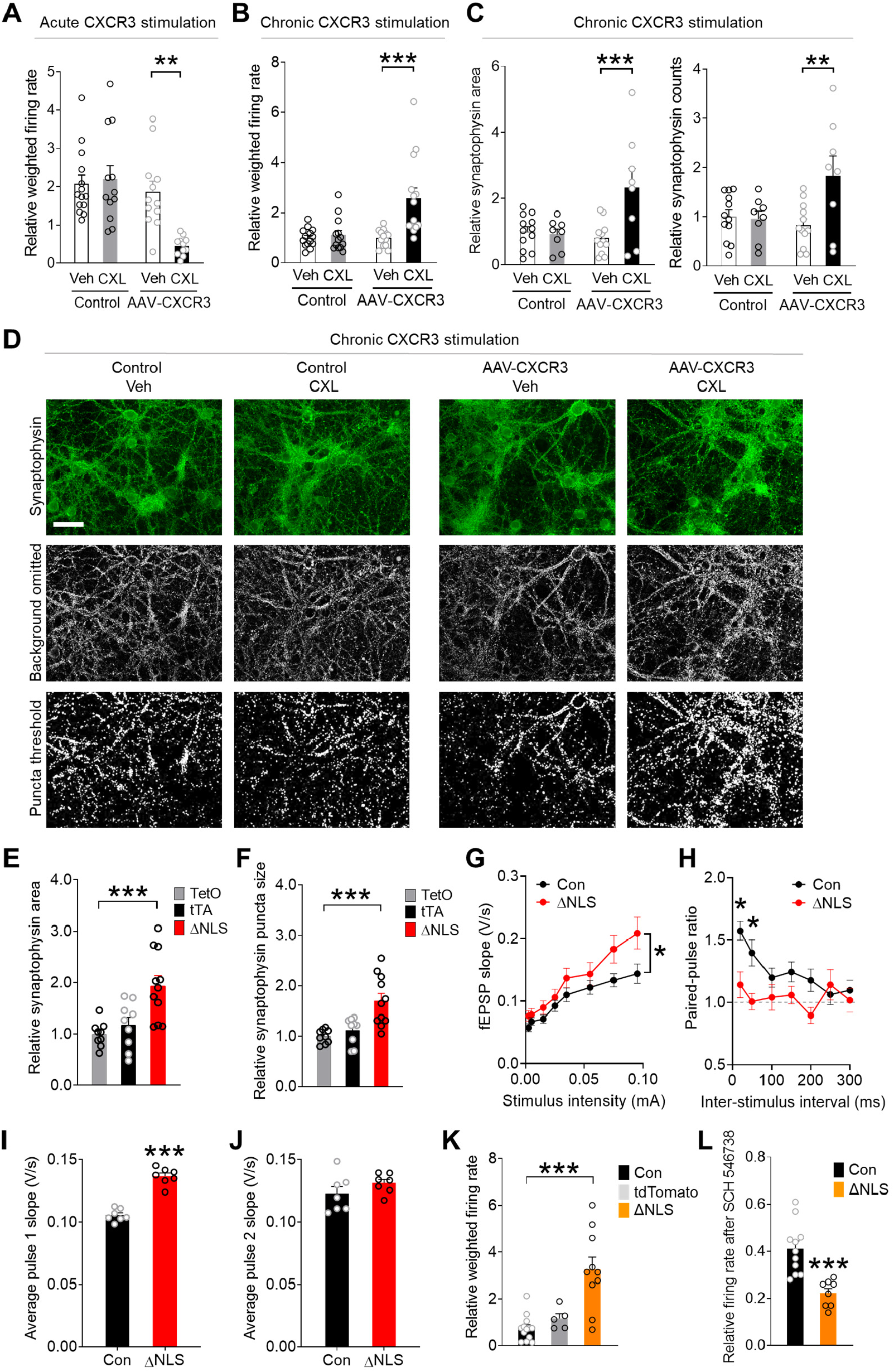
Astrocytic TDP-43 and neuronal CXCR3 promote hyperexcitability and aberrant presynaptic function. (**A** and **B**) Primary wild-type neurons were transduced with AAV PHP.eB *Syn*-CXCR3-HA-neurexin1α to target CXCR3 to presynaptic terminals. Control cultures were not transduced. (**A**) Recombinant CXCL11 (CXL, 200 nM) or vehicle (Veh) was applied acutely after a 30-min baseline recording using the MEA. Firing rates at 1 min post-stimulation were normalized to the average baseline rate per well. Two-way ANOVA: *F*(1, 45) = 8.38, p = 0.0058 for interaction effect; *F*(1, 45) = 5.83, p = 0.02 for CXL effect. Bonferroni post-hoc test: **p < 0.01 vs. Veh. n = 11–14 culture wells per genotype and treatment condition. (**B**) Recombinant CXCL11 (CXL, 200 nM) or vehicle (Veh) was applied chronically (24 h) after a 30-min baseline recording. Firing rates were normalized to the average baseline rate per well. (**C** and **D**) Quantification (C) and representative images (D) of immunoreactivity area and puncta counts for the presynaptic marker synaptophysin in primary wild-type neurons transduced with AAV PHP.eB *Syn*-CXCR3-HA-neurexin1α and treated chronically (3 days) with recombinant CXCL11 (CXL, 200 nM) or vehicle (Veh). Two-way ANOVA: *F*(1, 50) = 8.85, p = 0.0045 for interaction effect (firing rate); *F*(1, 50) = 11.73, p = 0.0012 for CXL effect (firing rate); *F*(1, 36) = 8.65, p = 0.0057 for interaction effect (area); *F*(1, 36) = 6.24, p = 0.017 for interaction effect (counts). Bonferroni post-hoc test: **p < 0.01, ***p < 0.001 vs. Veh. n = 5–14 culture wells per genotype and treatment condition. Scale bar: 50 µm. (**E** and **F**) Quantification of immunoreactivity for the presynaptic marker synaptophysin in the CA1 radiatum of singly transgenic controls and doubly transgenic ΔNLS mice. One-way ANOVA: *F*(2, 26) = 9.94, p = 0.0006 (E); *F*(2, 26) = 11.62, p = 0.0002 (F); Dunnett’s post-hoc test: ***p < 0.001 vs. TetO. n = 9–12 mice per genotype. (**G** and **H**) Recordings in acute hippocampal slices from singly transgenic controls (TetO and tTA; Con) and doubly transgenic ΔNLS male mice at 5–6 months of age. (**G**) Basal synaptic transmission at increasing stimulus intensities. Mixed-effects model: *F*(7, 344) = 2.10, p = 0.043 for interaction effect, *F*(1, 50) = 4.19, p = 0.046 for genotype effect. n = 28 recordings from 6 Con male mice and 24 recordings from 5 ΔNLS male mice. (**H**) Paired-pulse facilitation is shown as a ratio of fEPSPs in response to the second pulse as compared to the first pulse. Ratios are plotted as a function of interstimulus interval. Mixed-effects model: *F*(6, 338) = 2.43, p = 0.026 for interaction effect, *F*(1, 60) = 7.81, p = 0.007 for genotype effect. Bonferroni’s post-hoc test: *p < 0.05 vs. Con. n = 37 recordings from 6 Con male mice and 25 recordings from 5 ΔNLS male mice. (**I** and **J**) Average fEPSPs in response to the first (I) and second (J) stimulus independent of interval. Student’s *t* test: ***p < 0.001. (**K** and **L**) Primary wild-type neurons were co-cultured with hippocampal *Aldh1l1*-Cre astrocytes that were transduced with AAV PHP.eB-*hGfaABC*_*1*_*D*-DIO-hTDP43-ΔNLS (ΔNLS) or AAV PHP.eB pAAV-FLEX-tdTomato. Control cultures were not transduced. One-way ANOVA: *F*(2, 25) = 15.93, p < 0.0001 (K). Some wells received the selective CXCR3 blocker SCH 546738 (12 nM). Dunnett’s post-hoc test: ***p < 0.001 vs. Con. Student’s *t* test: ***p = 0.0003. n = 5–14 culture wells per genotype and treatment condition.

In contrast to these acute effects, chronic stimulation of CXCR3 increased spontaneous neuronal activity (**Fig. 9B**), indicating that CXCR3 also promotes long-lasting increases in neuronal firing, likely through chronic changes in the presynaptic compartment. Indeed, chronic stimulation of CXCR3 enhanced the levels of synaptophysin-positive puncta, a marker of presynaptic vesicles, but did not affect PSD-95, a marker of postsynaptic compartments (**Fig. 9C– D** and **fig. S7C**). Of note, neurons transduced with AAV-CXCR3 but not treated with chemokines, and neurons treated with chemokines but not transduced with AAV-CXCR3 performed similarly to untransduced vehicle-treated control neurons, ruling out nonspecific effects of the AAV vector and treatment on neuronal responses.

To test whether similar presynaptic changes were present in the hippocampus of ΔNLS mice, we compared the levels of different synaptic markers by quantitative immunofluorescence. Similar to neuronal cultures overexpressing presynaptic CXCR3, doubly transgenic ΔNLS mice had increased levels of synaptophysin-positive puncta (**Fig. 9E–F**) without marked changes in the levels of PSD-95 or gephyrin, which are markers of excitatory and inhibitory postsynaptic zones, respectively (**fig. S7D**–**E**). There were also no detectable changes in the levels of bassoon, a scaffolding protein in excitatory and inhibitory presynaptic compartments, and no changes in synaptotagmin-2, a marker of inhibitory presynaptic vesicles (*90, 91*) (**fig. S7F**–**G**). These results are consistent with recent findings that G_i/o_-coupled receptors can enhance the number of presynaptic vesicles without altering the number of synaptic zones (*92, 93*). Thus, astrocytic TDP-43 modulates excitatory presynaptic compartments without markedly changing the density of synaptic zones.

We next investigated whether astrocytic TDP-43 altered functional electrophysiological readouts of hippocampal transmission and presynaptic release. For these experiments, acute hippocampal slices were obtained from doubly transgenic ΔNLS mice or littermate controls at 5– 6 months of age. The Schaffer collateral pathway was stimulated while recordings of field excitatory postsynaptic potentials (fEPSPs) were obtained in the striatum radiatum of the dorsal CA1 region. We found that ΔNLS mice had increased basal synaptic transmission as evidenced by enhanced fEPSPs (**Fig. 9G**). This effect was most apparent at higher stimulus intensities. The mice also had impaired paired-pulse facilitation, as reflected by lower fEPSC_2_/fEPSC_1_ ratios (**Fig. 9H**). Notably, ΔNLS mice had increased responses to the first but not second pulses as compared to control mice (**Fig. 9I–J**), indicative of enhanced probability of neurotransmitter release, which likely contributed to the observed increases in basal synaptic transmission. No significant differences were detected in fiber volley amplitudes, which reflect presynaptic action potentials, or in linear regressions of fEPSP slopes and fiber volleys (**fig. S7H**–**I**).

To further test whether the presence of astrocytes with TDP-43 alterations is sufficient to affect neuronal activities and to confirm that this effect is not dependent on other cell types, we generated primary astrocytic-neuronal co-cultures in which neurons were derived from NTG mice and astrocytes were derived from transgenic *Aldh1l1*-Cre mice to enable astrocyte-selective transgene expression. Prior to the addition of neurons to the cultures, isolated *Aldh1l1*-Cre astrocytes were transduced with a PHP.eB AAV vector encoding Cre-dependent hTDP43-ΔNLS under the control of the *hGfaABC*_*1*_*D* promoter. Consistent with our findings in ΔNLS mice, NTG neurons co-cultured with *Aldh1l1*-Cre astrocytes expressing hTDP43-ΔNLS had altered spontaneous firing patterns as compared to neurons co-cultured with control *Aldh1l1*-Cre astrocytes (**Fig. 9K**). Notably, spontaneous neuronal activities were suppressed by selective blockade of CXCR3 with SCH 546738 (12 nM) to approximately 40% of baseline activities, and this suppressive effect was significantly more pronounced in neurons cultured in the presence of hTDP43-ΔNLS-expressing astrocytes (**Fig. 9L**), revealing an increased involvement of CXCR3. Together with our findings in isolated neurons and hippocampal slices, these results suggest that astrocytic TDP-43 is linked to CXCR3-dependent neuronal hyperexcitability.

Given these findings, astrocytic TDP-43 alterations may promote memory loss in part through CXCR3-induced effects on hippocampal function. Thus, we next tested whether genetic ablation of the gene encoding CXCR3 can prevent TDP-43-related memory loss. For these experiments, we generated *Aldh1l1*-Cre mice that were either wild-type (WT) or functional null (knockout, KO) for *Cxcr3* and performed bilateral injections of the PHP.eB AAV vector encoding hTDP43-ΔNLS into the hippocampal formation at 4–9 months of age (**Fig. 10A–B**). Given that *Cxcr3* is X-linked, we used littermate males that were either WT or KO for *Cxcr3* and expressed Cre recombinase, which enabled cell type-selective hTDP-43 expression in hippocampal astrocytes. The mice were then assessed in the Morris water maze at 7–12 months of age.

**Fig. 10.**
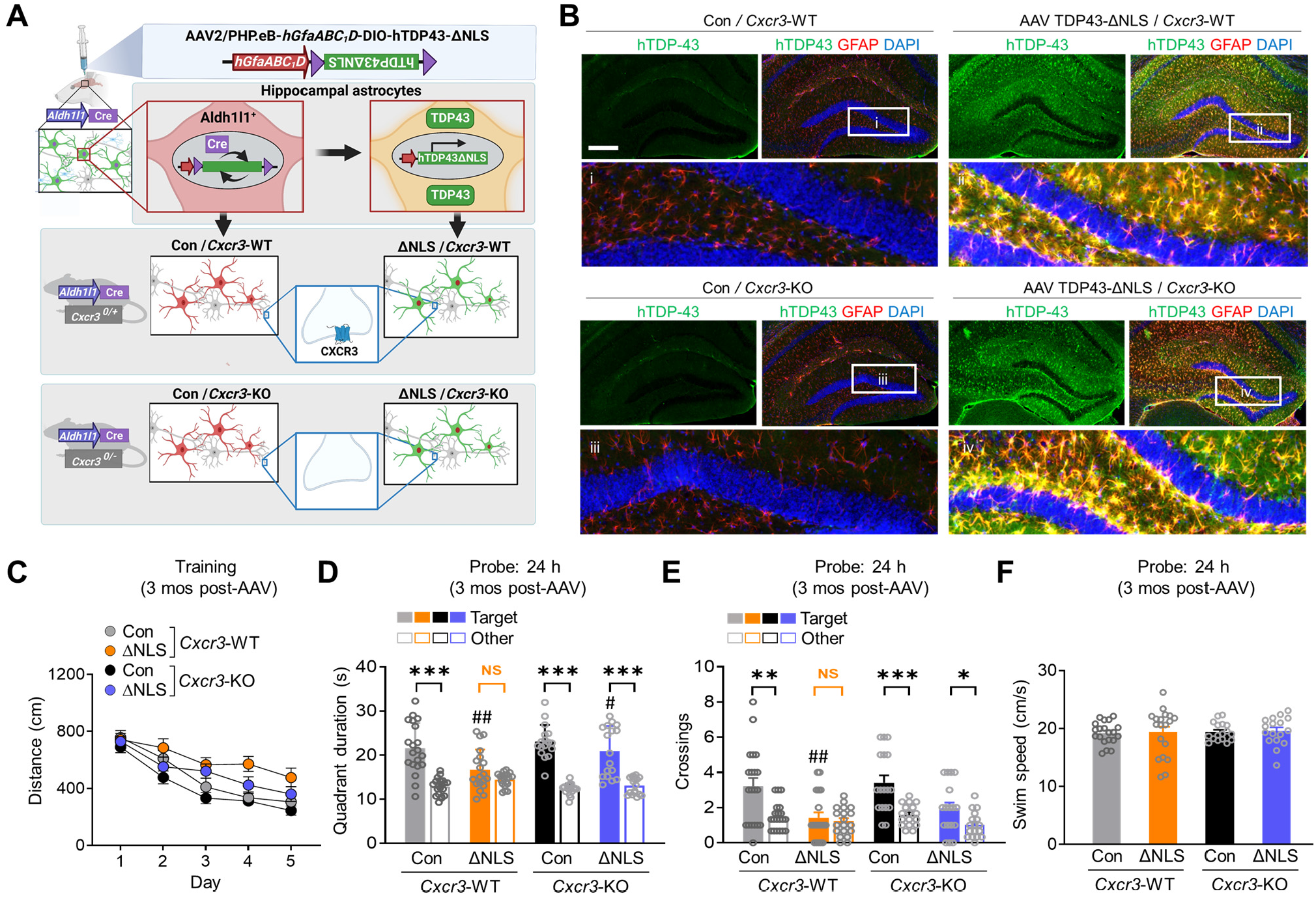
Ablation of CXCR3 alleviates astrocytic TDP-43-linked memory deficits. (**A**) Schematic of the experimental design. Transgenic *Cxcr3*-WT or *Cxcr3*-KO male mice on a *Aldh1l1*-Cre background were injected with AAV PHP.eB-*hGfaABC*_*1*_*D*-DIO-hTDP43-ΔNLS (ΔNLS) or saline (Con) at 4–9 months of age and tested in the Morris water maze at 7–12 months of age. Control AAV injections in NTG mice are shown in Figure 4 and Extended Data Fig. 4. (**B**) Representative images of human-specific TDP-43 (green) and the astrocytic marker GFAP (red). DAPI (blue) was used to visualize nuclei. Yellow indicates overlay of green and red channels. Insets i–vi show magnified views. Scale bar: 300 µm. (**C**) Distance traveled to reach the platform during hidden platform training (four trials per session, one session per day). Mixed-effects model: *F*(1, 70) = 17.84, p < 0.0001 for ΔNLS effect, *F*(1, 70) = 8.93, p = 0.0039 for *Cxcr3*-KO effect, *F*(1, 70) = 0.12, p = 0.73 for ΔNLS x *Cxcr3*-KO interaction effect. n = 20 Con / *Cxcr3-*WT, 19 ΔNLS */ Cxcr3*-WT, 17 Con / *Cxcr3*-KO, and 17 ΔNLS / *Cxcr3*-KO mice. (**D** and **E**) Probe trial conducted 24 h after training. Durations in target and non-target (Other) quadrants. Two-way ANOVA (Target): *F*(1, 69) = 8.82, p = 0.004 for ΔNLS effect, *F*(1, 69) = 5.94, p = 0.017 for *Cxcr3*-KO effect (D); *F*(1, 69) = 16.3, p = 0.0001 for ΔNLS effect (E). Bonferroni’s post-test: ##p < 0.01 vs. Con / *Cxcr3-*WT, #p < 0.05 vs. ΔNLS / *Cxcr3-*WT. Student’s *t* test with Welch’s correction: ***p < 0.001, **p < 0.01, *p < 0.05 vs. Other. No significant preference for target (NS). (**F**) Mice were assessed for swim speeds in the 24 h probe trial.

Similar to other results (**Fig. 4**), *Aldh1l1*-Cre/*Cxcr3*-WT mice that received AAV encoding hTDP43-ΔNLS had impaired memory as compared to control mice that lacked hTDP43-ΔNLS (**Fig. 10C–E**), without changes in swim speeds (**Fig. 10F**). In contrast, *Aldh1l1*-Cre/*Cxcr3*-KO mice with hTDP43-ΔNLS performed more similarly to control mice lacking hTDP43-ΔNLS (**Fig. 10D–E**), suggesting that alterations in astrocytic TDP-43 cause memory loss in a CXCR3-dependent manner. Also, in control conditions, the behavior of *Aldh1l1*-Cre*/Cxcr3*-KO mice was not different from *Aldh1l1*-Cre*/Cxcr3*-WT mice, suggesting that CXCR3 impairs memory upon changes in astrocytic function. Altogether, our study reveals that astrocytic TDP-43 is linked to maladaptive antiviral changes and increased chemokine signaling that disrupts hippocampal synaptic transmission and contributes to memory loss (**fig. S8**).

## Discussion

Memory loss is common in aging-related neurological disorders, including AD, ALS-FTD, hippocampal sclerosis, and other conditions. However, the exact causes of memory loss are not clear and treatment options are limited. Increasing evidence implicates glial dysfunction and abnormal glial-neuronal interactions in various pathophysiological processes (*37-39*). In particular, astrocytes have been implicated in various CNS disorders (*94-98*) and, like neurons, are functionally diverse (*99-101*) and affect information processing (*102-104*). Aberrant changes in astrocytes can contribute to behavioral and cognitive deficits and promote memory loss associated with aging and disease (*42, 105-108*). However, the mechanisms by which astrocytes impair memory and other cognitive processes are not fully defined.

Here, we demonstrate that human hippocampal astrocytes accumulate extranuclear TDP-43 in AD and FTD. Astrocytic TDP-43 accumulation in the hippocampus was sufficient to impair memory, but not other neurocognitive functions, and it altered hippocampal neural activity and presynaptic function. Consistent with the selective impairments in memory, we detected marked increases in interferon-inducible chemokines preferentially in hippocampal astrocytes and increases in the corresponding chemokine receptor CXCR3 in hippocampal presynaptic terminals. These findings suggest that astrocytes in the hippocampus have a distinct response to TDP-43 alterations as compared to astrocytes in other brain regions. Indeed, widespread and chronic transgene expression in astrocytes did not cause motor impairments, early mortality, or other severe deficits. In contrast, animal models with analogous TDP-43 manipulations in neurons have severe ALS-associated phenotypes, including motor impairments and early mortality (*19, 109*). These distinct and selective effects suggest that responses to TDP-43 accumulation are cell type-specific and heterogeneous across astrocytes in different brain regions.

Similar to TDP-43, ALS-linked mutations in superoxide dismutase (SOD1) cause neurotoxicity, motor impairments, and early mortality. Analogous to our findings, astrocyte-targeted expression of mutant SOD1 is not sufficient to trigger onset of motor neuron disease in mice (*110*). However, astrocytic SOD1 is required for disease progression, and astrocytes carrying mutant SOD1 are selectively damaging to isolated motor neurons but not other neuronal subtypes (*111, 112*). Similarly, AD-linked tau accumulation in hippocampal astrocytes promotes selective neuronal deficits (*113*). Altogether, our findings and previous work indicate that various dementia-linked protein alterations in astrocytes cause highly context-dependent effects on neurons and might facilitate selective neuronal vulnerability that contributes to variable disease manifestations in neurodegenerative conditions.

Most dementia cases involve dysregulation of wild-type rather than mutant TDP-43. Overexpression of wild-type TDP-43 in model systems is sufficient to impair cell function at least in part through the effects of TDP-43 outside the nucleus (*57*). We found that expression of either hTDP43-WT or hTDP43-ΔNLS, but not control proteins targeted to hippocampal astrocytes induced progressive memory deficits, suggesting that even modest alterations in wild-type TDP-43 within hippocampal astrocytes can impair memory and contribute to dementia-associated cognitive decline. Hippocampal astrocytes were not similarly vulnerable to control vectors, thus ruling out nonspecific effects of AAV injections or chronic protein overexpression as major drivers of the observed phenotype. Of note, microglia and other neural cells in the hippocampus might indirectly modulate astrocytic functions and the effects of TDP-43 alterations. Indeed, microglial progranulin and TREM2 insufficiency may contribute to TDP-43 pathology (*58, 114*).

Astrocytic TDP-43 alterations were accompanied by cell-autonomous changes in antiviral gene expression, increased phosphorylation of NF-κB and STAT3, and functional changes in astrocytic innate defense against viral pathogens. Together, these results point to an aberrant TDP-43-linked antiviral phenotype (aTAP) that may impact cognitive function as well as neuroimmune responses. Our study focused on astrocytes, but TDP-43 is present in most cell types and increasingly linked to interferon-related pathways in different cell populations (*115-117*), suggesting that aTAP is not specific to astrocytes. Indeed, TDP-43 accumulation in neurons triggers the antiviral cGAS-STING pathway, at least partly through abnormal mitochondrial DNA release (*57*). We did not detect changes in astrocytic cGAS or STING genes, possibly because aTAP engages distinct mechanisms in different cell types. Indeed, antiviral signaling involves multiple dynamic and cell-specific mechanisms (*118*). TDP-43 can also affect retrotransposon activity (*119, 120*), which may also contribute to aTAP. Thus, TDP-43 likely influences multiple intracellular targets that affect neuroimmune signaling and antiviral cascades.

Although we focused primarily on the neurocognitive effects, our results in isolated cells implicate TDP-43 in modulating innate responses to viral pathogens. A link between dementia and viral infections has been suggested (*121-123*), but the effects of TDP-43 pathology on neural responses to infections are not known. We tested three different viral pathogens, used several independent methods to measure viral infections, and assessed different time-points and viral doses. Convergent results across these different conditions suggest that alterations in TDP-43 allow pathogens to exploit weaknesses in astrocytic antiviral responses, which might affect innate immunity in the brain.

Interferon-related pathways in astrocytes and other neural cells modulate brain function and have been implicated in AD, ALS-FTD, and other CNS disorders (*71, 73, 76, 77, 124-127*), but the roles of TDP-43 in these pathways have not been fully elucidated. We found that alterations in TDP-43 increase astrocytic interferon-inducible chemokines, among other genes, and promote presynaptic increases in neuronal CXCR3, the shared receptor that is likely overactivated by the increased levels of chemokines. Although previous studies have reported CXCR3 expression in microglia (*128*) and infiltrating immune cells (*129*), we did not detect increases in CXCR3 in non-neuronal areas within the hippocampus. Similar to our results, CXCR3 has been detected in human neurons and neuronal processes (*67*). Acute activation of CXCR3 suppressed neuronal activity whereas chronic activation of CXCR3 increased neuronal activity. In a similar manner, neurons maintained in the presence of astrocytes with chronic TDP-43 accumulation had increased spontaneous neuronal activity, which was reduced by pharmacological inhibition of CXCR3. These results suggest that alterations in astrocytic TDP-43 promote CXCR3-dependent neuronal hyperexcitability. Furthermore, ΔNLS mice had increases in hippocampal transmission and presynaptic release probability, which may promote an abnormal shift to low-pass filtering of excitatory firing and thereby impair the encoding of spatial memory within the hippocampus. In support, genetic ablation of CXCR3 alleviated memory deficits but did not affect memory in mice without astrocytic manipulation. Thus, alterations in astrocytic TDP-43 cause neural hyperexcitability and memory deficits that are at least partly dependent on CXCR3.

These findings implicate blockers of chemokines and CXCR3 as novel therapeutic approaches for TDP-43-associated cognitive impairments. Notably, human CXCL10 levels are increased in progranulin-linked FTD (*86*), AD (*130*), and amnestic mild cognitive impairment (*130*), either in the cerebrospinal fluid or in astrocytes (*67*), and these levels correlate with cognitive performance (*130*). Moreover, neural hyperexcitability has been reported in dementia (*131, 132*) and global knockout of CXCR3 prevents memory deficits in transgenic mice with amyloid-β pathology (*133*). CXCR3 blockers have reached clinical trials for peripheral inflammatory conditions (*134*), but have not yet been tested in patients with dementia or other cognitive disorders.

In summary, our findings suggest that TDP-43 alterations in astrocytes contribute to cognitive decline in dementia. We describe a novel chemokine-mediated mechanism that is likely downstream of astrocytic TDP-43-linked antiviral changes that affects hippocampal presynaptic function and neuronal activities. Together, our results implicate astrocytic TDP-43 impairments and aTAP in dementia pathogenesis and point to chemokine signaling as a potential therapeutic target.

## Materials and Methods

### Mice

All animal experiments were conducted in accordance with guidelines set by the Institutional Animal Care and Use Committee of Weill Cornell Medicine. Mice were housed in groups of 2–5 mice per cage and maintained on a 12-h light/dark cycle with *ad libitum* access to food and water. Experiments were conducted during the light cycle and included littermate controls. Regulatable and astrocyte-specific expression of human TDP-43 was achieved using transgenic *hGFAP*-tTA mice with a tet-off tetracycline transactivator (tTA) element downstream of the human glial fibrillary acidic protein (*hGFAP*) promoter (kindly provided by Dr. Lennart Mucke, Gladstone Institutes, San Francisco, CA) (*42, 135*). *hGFAP*-tTA mice (B6.Cg-Tg(GFAP-tTA)110Pop/J; Jackson Laboratory strain #005964) were crossed with *tetO*-hTDP43-ΔNLS mice (B6;C3-Tg(tetO-TARDBP*)4Vle/J; Jackson Laboratory strain #014650), which contained a tet operator (tetO) upstream of the human TARDBP gene with a mutated nuclear localization sequence (*tetO*-hTDP43-ΔNLS) leading to expression of cytoplasmic human TDP-43 (*19*). In doubly transgenic mice (referred to as ΔNLS mice), doxycycline (DOX) can bind to tTA to prevent *tetO*-mediated transgene expression. DOX-supplemented chow (200 mg/kg, VWR, 89067-462) was provided to breeding pairs and offspring until weaning (P21) to prevent expression of transgene during embryonic and postnatal development. Thereafter, mice were maintained on standard laboratory chow (Purina 5053) to allow the induction of transgene expression. Because the parent *tetO*-hTDP43-ΔNLS strain was on a B6/C3 hybrid background, we backcrossed this line onto the C57Bl/6J background (Jackson Laboratory strain #000664) for five generations.

Aldehyde dehydrogenase 1 family, member L1 (*Aldh1l1*)-Cre transgenic mice (B6;FVB-Tg(*Aldh1l1*-cre)JD1884Htz/J) were obtained from the Jackson Laboratory (strain #023748) and backcrossed onto the C57Bl/6J background. *Aldh1l1*-Cre mice express Cre recombinase downstream of the astrocytic *Aldh1l1* promoter. Homozygous *Aldh1l1*-Cre mice were crossed with transgenic mice null for the chemokine receptor CXCR3 (B6.129P2-Cxcr3tm1Dgen/J; Jackson Laboratory strain #005796) to create *Aldh1l1*-Cre/*Cxcr3*-WT and *Aldh1l1*-Cre/*Cxcr3*-KO male mice. Hippocampal tissue from transgenic mice expressing mutant human tau-P301S (B6;C3-Tg(Prnp-MAPT*P301S)PS19Vle/J, Jackson Laboratory stock #008169) was used for gene expression comparisons.

### Adeno-associated virus (AAV) preparation

Wild-type and mutant forms of human TDP-43 were cloned into plasmids for pAAV-mediated, astrocyte-specific expression in two stages. First, the truncated human astrocyte-specific promoter *hGfaABC*_*1*_*D* (*54*) was digested from pAAV-GFAP-EGFP (donated by Dr. Bryan Roth, Addgene Plasmid #50473; RRID #Addgene_50473) and cloned into pAAV-EF1a-DIO-hM4D(Gi)-mCherry (donated by Dr. Bryan Roth, Addgene Plasmid #50461; RRID #Addgene_5046) in place of the EF1a promoter using Anza 28 MluI and Anza 14 SalI (ThermoFisher). The inverted hM4D(Gi)-mCherry coding sequence in the resulting vector was replaced with inverted human wild-type or NLS1 TDP-43 coding sequences (*136*) by PCR amplification. Primers were designed to contain NheI and SgsI restriction sites 5’ and 3’ to the TDP-43 coding sequence, respectively (NheI primer = 5’-TGT CGC TAG CGC CAC CAT GTC TGA ATA TAT TCG G-3’; SgsI primer = 5’-AAG GCG CGC CCT ACA TTC CCC AGC CAG AAG-3’). Amplicons were digested and gel-purified before ligation into similarly prepared pAAV-*GfaABC*_*1*_*D*-DIO backbone. The pcDNA3.2 TDP-43 YFP vectors were donated by Aaron Gitler (Addgene plasmids # 84911, 84912, RRID: Addgene_84911, Addgene_84912).

Mouse CXCR3 with a C-terminal HA tag was targeted to neuronal presynaptic terminals using the neurexin-1α targeting sequence, as described (*89*). A gBlock gene fragment (Integrated DNA Technologies) encoding mCherry-T2A-CXCR3-2HA-neurexin1α (axon targeting sequence) was designed following sequences made available by Scott Sternson (Addgene plasmid #52523; RRID #Addgene_52523), synthesized and cloned into an AAV-hSyn1 expression vector (Addgene plasmid #50458; RRID #Addgene_50458; donated by Dr. Bryan Roth) using SalI and EcoRI restriction digest to generate pAAV-*hSyn1*:mCherry-T2A-CXCR3-2HA-nrxn1α. NEB 5-alpha cells (New England Biolabs) were transformed with pAAV constructs and the integrity of inverted terminal repeats and expression-related elements in selected clones were confirmed by sequencing and restriction digests. pAAV2/PHP.eB or pAAV2/DJ particles were produced by the Stanford University Neuroscience Gene Vector and Virus Core or the University of Pennsylvania Vector Core. PHP.eB capsid vectors were provided courtesy of Dr. Viviana Gradinaru and Dr. Benjamin Deverman at the California Institute of Technology. PHP capsids are a modification of AAV9 provided by the University of Pennsylvania (*55, 137*).

### Surgery and AAV microinjections

Mice were anesthetized with sterile Avertin (2,2,2-tribromoethanol, 400–500 mg/kg body weight, Acros Organics) and the hair was removed from the surgical area. Mice were secured in a stereotaxic frame (Kopf Instruments) and 1 mm-diameter openings were made bilaterally in the skull using a mounted drill (Kopf Instruments). Meloxicam (2 mg/kg) was injected subcutaneously, and bupivacaine (1 mg/kg) was applied topically to relieve pain. Stereotaxic coordinates used for hippocampal injections were (from bregma) anterior/posterior: -2.1, medial/lateral: -/+ 1.7, and dorsoventral: -2.0 (for mice under 23 g body weight) or -2.1 (for mice over 23 g body weight). A blunt 32-gauge, 0.5-inch-long needle attached to a 5 µl Hamilton syringe was mounted to the stereotaxic frame and controlled using a Micro 4 Microsyringe Pump (World Precision Instruments) to infuse 0.5 µl AAV2/PHP.eB-*hGfaABC*_*1*_*D*-DIO-hTDP43-WT (1.2 × 10^13^ particles/µl), AAV2/PHP.eB-*hGfaABC*_*1*_*D*-DIO-hTDP43-ΔNLS (1.3 × 10^13^ particles/µl), or AAV2/PHP.eB-*hGfaABC*_*1*_*D*-DIO-hM4Di-mCherry (3.15 × 10^12^ particles/µl) at right and left injection sites at a rate of 0.1 µl/min, after which the needle was left in place for an additional 5 min. After needle withdrawal, the surgical site was sealed with Vetbond tissue adhesive (3M). Mice were monitored under a heating lamp until fully recovered and returned to their home cage.

### Behavioral testing

Experimental groups were distributed randomly across home cages and consisted of age-matched littermates of both sexes. Experimenters were blinded to genotypes and mice were tested in random order. Before most behavioral testing, except for the elevated plus maze, all mice were handled for approximately 2 min per day for 7 days. Mice that were injured or in poor health, independent of genotype, were excluded from behavioral testing. Tests were performed under white light, unless otherwise noted. For all test days, mice were acclimated to the testing room for 1 h prior to testing.

#### Elevated plus maze

The plus-shaped maze consisted of two enclosed arms and two open arms elevated 60–70 cm above the ground. Furthermore, tape was attached to the ends of the open arms (5 cm from the end of the arm) to limit falls. After 1 h of habituation, mice were placed at the center of the maze facing an open arm. Mice could freely explore the four arms for 5 min. Time and distance traveled in each arm and center area were video recorded and tracked using EthoVision XT video tracking software (Noldus Information Technology Inc. Leesburg, VA). The apparatus was cleaned with 70% alcohol between mice.

#### Open field test

Mice were placed in the center of a clear plastic chamber (41 × 41 × 30 cm) with two 16 × 16 photobeam arrays detecting horizontal and vertical movements. To measure context-dependent habituation in the open field, the chambers were surrounded by distinct cues that were maintained across test days. Mice were acclimated to the chamber in 2 × 5-min trials with a 3-h inter-trial interval and assessed in the same chambers 1 and 14 days after habituation. Light in the room was set to 75% red light to limit the anxiolytic effect of 100% white light. Total exploration, rearing and percent time spent in the center of the arena were measured with an automated Flex-Field/Open Field Photobeam Activity System (San Diego Instruments, San Diego, CA). The apparatus was cleaned with 70% alcohol between mice.

#### Rotarod

Mice were placed on the Rotarod (Rotamex-5 0254-2002L) that was suspended 25.5 cm from a soft surface. The speed of rotation was either held constant at 12 RPM or increased from 4 to 40 RPM gradually at an acceleration rate of 0.3 RPM/s. Mice were tested on the rod in 3 trials with approximately 30 min inter-trial intervals. The latency to fall off the rod was recorded and reported as the average of three trials. Equipment was cleaned with 70% ethanol between trials and the light in the room was set to red light.

#### Morris water maze

The maze consisted of a 122-cm-diameter pool filled with water (20 ± 2°C) made opaque with nontoxic white tempera paint (Colorations powder tempera paint). Spatial cues were set up around the pool prior to testing. All mice underwent one session of 3–4 pre-training trials in which they swam in a rectangular channel (15 cm × 122 cm) with a square platform (14 × 14 cm) hidden 0.5 cm below the water surface in the middle of the channel. If a mouse did not reach the platform within 10 s, it was guided onto the platform by the experimenter and remained on the platform for 10 s before it was returned to its cage. One to three days following pre-training, mice underwent hidden platform training in the circular water maze.

For hidden platform training, the platform was submerged 1.5 cm below the surface. All mice underwent one session of 4 trials for 3–5 consecutive training days. For each trial, the platform location remained the same, but the mice were dropped in 4 different locations. The maximum time allowed per trial was 60 s. If a mouse did not find or mount the platform, it was guided to the platform by the experimenter. All mice were allowed to sit on the platform for 10 s after each training trial.

Probe trials were performed 24 h and 72 h after the last hidden platform-training day. For probe trials, the platform was removed, and mice were allowed to swim for up to 60 s per trial. The drop location for the probe trials was 180° from the platform location used during hidden platform training. After 60 s, mice were guided to the platform location before removal from the pool and returned to its cage.

If the mice had any problems in learning where the platform is located, we performed a cued platform training 24 h after probe testing. All mice underwent one session of 4 trials of the cued platform training. The cued (visible) platform training was performed using a new platform location and a clearly visible cue (a colorful 15-cm pole on top of the platform). All behavior was recorded and analyzed with an Ethovision XT video tracking system (Noldus). Escape latencies, distance traveled, swim paths, swim speeds, platform crossings and proximity to the platform were recorded automatically for subsequent analysis.

#### Novel object recognition test

Mice were habituated to the testing chamber (40 × 40 cm) for 15 min. A day after habituation, mice were exposed to two identical objects in the same chamber and allowed to explore freely for 10 min once per day for two consecutive days. The next day, mice were presented with one object used during training and one unfamiliar (novel) object of a different shape and texture in the same chamber, and the mice were allowed to explore for 15 min during a test trial. The objects used for training and testing were assigned randomly to each mouse to avoid object bias and which of the familiar objects was replaced with a novel object was varied randomly between mice to control for location bias. Chamber and objects were cleaned with Clidox-S (Pharmacal; 1:18:1 dilution) after each mouse. Behavior was recorded and analyzed with an Ethovision XT video tracking system (Noldus) the time that the mice spent next to each object were scored.

#### Social interaction test

Mice were allowed to freely explore a three-chamber arena (side chambers were 22.86 cm x 42.2 cm; middle chamber was 21.59 cm x 42.2 cm) with two empty inverted wire cups (8.5 cm diameter) in each side chamber. Chambers were divided by clear plexiglass dividers, each with a half-circular opening at the bottom to serve as a free passageway between chambers. After 10 min of exploration by a test mouse, a novel mouse of the same sex as the test mouse was placed under one inverted wire cup and the other wire cup was left empty. The test mouse was allowed to freely explore the three-chamber arena for another 10 min to explore the novel partner. Each novel mouse used for testing was assigned randomly to each side to avoid location bias. Chambers and wire cups were cleaned with 70% ethanol after each mouse. Behavior was recorded and analyzed with the Ethovision XT video tracking system (Noldus). The total time that the test mouse spent next to each inverted wire cup with a mouse or empty wire cup was scored. The light in the room was set to red light.

#### Marble-burying test

Mice were placed in large cages (12 cm x 12 cm x 7.25 cm) covered with mouse bedding material to a depth of 5 cm. During each trial, 20 standard glass black marbles were gently placed on the surface of the bedding in a grid pattern. A mouse was placed in the center of the cage and allowed to explore for 30 min. A marble was scored as buried when it was at least 3/4 covered with bedding. The light in the room was set to red light. Between trials, the bedding was changed, and the cages and marbles were cleaned with 70% ethanol.

#### Nestlet-shredding test

Mice were single-housed with one cotton fiber nestlet (5 cm x 5 cm, 5 mm thick, ∼2.5 g each) for 60–90 min. Nestlets were weighed before putting them in the cages. After returning the mouse to its home cage, whole nestlet pieces (not the shredded nestlet) were removed, dried overnight, and weighed.

#### Grooming test

Mice were sprayed three times with a light water mist on the back and neck areas. The mice were placed in large cages (12 cm x 12 cm x 7.25 cm) with no bedding. Behavior was recorded with the Ethovision XT video tracking system (Noldus) for 10 min. Cages were cleaned with 70% ethanol after each mouse. The amount of time spent grooming the head and the rest of the body was recorded and analyzed. The light in the room was set to red light.

#### Pole test

A vertical metal pole (1 cm diameter, 50 cm tall) with a heavy metal base was placed in the center of a clean cage covered with bedding. Individual mice were placed on the upper portion of the pole facing the ceiling for three trials. The amount of time spent turning at the top of the pole and climbing down are video recorded (Ethovision XT video tracking system, Noldus) and quantified. The apparatus was cleaned with 70% ethanol after each mouse. The light in the room was set to red light.

#### Wire hanging test

Individual mice were placed on top of a 2 mm thick metal cloth hanger that is securely attached 40 cm above the home cage containing extra padding. For three consecutive days the mouse was allowed to grasp the wire with the two forepaws for a maximum of 300 s for three trials. The latency of the mouse to full was recorded and quantified. The apparatus was cleaned with 70% ethanol after each mouse. The light in the room was set to red light.

### Cell culture experiments

#### Primary astrocyte cultures

All cultures were maintained at 37°C in a humidified 5% CO_2_-containing atmosphere. Cortices and/or hippocampi from wild-type (C57Bl/6J, Jackson Laboratory strain #000664), *Aldh1l1*-Cre (B6;FVB-Tg(*Aldh1l1*-cre)JD1884Htz/J, Jackson Laboratory strain #023748) or hemizygous doubly transgenic ΔNLS pups at postnatal day 1–3 were dissected in cold PBS to remove meninges and dissociated by manual trituration with a P1000 pipette in 1 ml fresh culture media consisting of high-glucose DMEM (Corning), 20% heat-inactivated FBS (VWR #89510-188), 1X GlutaMAX (ThermoFisher #35050061), and 1 mM sodium pyruvate (Thermo Scientific). In some experiments, culture media for isolated astrocytes derived from doubly transgenic TDP43-ΔNLS mice contained tetracycline-depleted FBS (VWR #97065-310) that was heat-inactivated for 30 min at 56°C in a water bath. To prevent transgene expression, some cultures were treated with 2 µg/µl doxycycline hyclate (Millipore-Sigma #D9891). Cell suspensions were diluted to 10 ml with media, filtered through a 70 μm cell strainer (VWR), centrifuged for 5 min at 300 g at 22°C, resuspended with culture media, and plated into cell culture dishes pre-coated with poly-D-lysine (75–150 kDa, 0.01% in water, filtered; Sigma #P6407 or MP Biomedical #0215017580). At DIV 4–5, the cells were washed to remove debris and given fresh media.

Wild-type and doubly transgenic astrocyte cultures were determined to contain 98.8% ± 0.4% (SE) astrocytes and 1.2% ± 0.4% (SE) microglia, based on immunolabeling for GFAP, Iba1, NeuN, and DAPI. These analyses were performed using a Nikon Eclipse Ti-S microscope with a Nikon PlanFluor 10X objective and NIS-Elements BR v5.02.01 acquisition software.

In doubly transgenic TDP43-ΔNLS cultures without doxycycline treatment, hTDP-43 expression was detected in 47.8 ± 4.7% (SE) of astrocytes, based on immunolabeling for hTDP-43 and the astrocyte marker GFAP. For these analyses, a stringent lower cutoff was determined using mean signal intensity in parallel cultures with doxycycline treatment, which accounted for nonspecific immunoreactivity and background fluorescence. For analyses of hTDP-43 distribution in cultures, images were captured with an LSM880 confocal microscope (Zeiss) equipped with a 63X objective (Zeiss) and Zen Black v2.3 SP1 FP3 acquisition software (Zeiss). Images were processed in FIJI v2.1.0/1.53c by subtracting the average background fluorescence and analyzing individual cells that had DAPI-positive nuclei and GFAP-positive cell bodies for levels of nuclear and extranuclear hTDP-43 immunoreactivity, respectively.

#### Primary neuronal cultures

Cortical and hippocampal neurons from postnatal day 0–1 wild-type or knockout mouse pups (B6.129P2-Cxcr3tm1Dgen/J; Jackson Laboratory strain #005796) were obtained as described previously (*138*), with minor modifications. Briefly, papain-dissociated cells were filtered through a 0.4 µm cell strainer (Corning #431750) to enrich for neurons, centrifuged at 500 g for 5 min to eliminate small debris, and suspended in complete primary neuronal medium consisting of B-27 Plus Neuronal Culture system (ThermoFisher #A3653401) and 1X GlutaMAX (ThermoFisher #35050061) without antibiotics. Cells were seeded at 50,000– 150,000 live cells per cm^2^ into plates coated with poly-D-lysine (0.01% w/v; Sigma P6407). Media was fully exchanged one day after plating (DIV 1) with subsequent half-media exchanges every 3–4 days.

#### Chemokine treatments in primary neurons

For immunostaining, primary wild-type neurons were cultured on poly-D-lysine-coated black walled µCLEAR 96-well plates (Greiner Bio-One #655090). For recording neuronal activity, primary wild-type neurons were plated onto poly-D-lysine coated 48-well CytoView multielectrode array (MEA) plates (Axion BioSystems) as described above. Neurons were transduced at DIV8 with 2×10^8^ AAV2/DJ-*hSyn1*:mCherry-T2A-Cxcr3-2HA-nrxn1a particles per well. Starting at DIV9, neuronal activity was recorded for 15– 30 min daily before and after treatment with recombinant CXCL11 (200 nM, BioLegend #573606) using the Maestro Pro MEA System (Axion BioSystems). Neuronal firing rates were analyzed using Neural Metric Software (Axion BioSystems).

For RT-qPCR or western blotting, primary wild-type neurons were cultured on TC-treated 24-well plates (Greiner Bio-One #662160). For chronic treatment with chemokines, neurons were treated with 200 nM CXCL11 (Biolegend #573606) or PBS vehicle (VWR #76018-870) starting at DIV4 with reapplication at 2X concentrations during half-volume feedings at DIV 7 and DIV 11. At DIV 14, neurons were fixed for immunostaining or harvested for RT-qPCR, as described in *Immunocytochemistry* or *Microfluidic qPCR*. To confirm the effects of chemokine stimulation on intracellular signaling, Neuro-2a cells (ATCC, #CCL-131) maintained in high-glucose DMEM (Corning), 10% heat-inactivated FBS (VWR #89510-188), 1X GlutaMAX (ThermoFisher #35050061), and 1 mM sodium pyruvate (Thermo Scientific), were transfected with AAV2/DJ-*hSyn1*:mCherry-T2A-Cxcr3-2HA-nrxn1a using Lipofectamine 3000. To assess the chronic effects of chemokine stimulation on synaptic markers, neurons were treated at DIV 10 with 200 nM recombinant mouse CXCL11 or PBS (vehicle) for 72 h before fixing with 4% PFA in 4% sucrose in PBS and immunostaining at DIV 13. For measuring G_i_-coupled phospho-signaling, after overnight starvation, Neuro2a were acutely treated with PBS (vehicle) or CXCL11 for 0, 2 or 10 min before harvesting as described in *Western blotting*.

#### Primary astrocytic-neuronal co-cultures

Cortical and/or hippocampal astrocytes at postnatal day 1–3 were cultured as described above. Prior to seeding neurons, near-confluent monolayers (typically 5–8 days after plating) were briefly rinsed of serum-containing medium. Neuronal suspensions were obtained from cortical and hippocampal tissue of postnatal day 0 mouse pups as described above and seeded at 50,000 live cells per MEA well atop existing rinsed astrocyte monolayers in neuronal media, as described above.

Astrocytes were plated onto poly-D-lysine-coated 48-well CytoView multielectrode array (MEA) plates (Axion BioSystems), as described above. After 6–8 days, cells were washed and transduced with 2 µl/well of AAV2/PHP.eB-*hGfaABC*_*1*_*D*-DIO-TDP43-WT (1.2×10^13^ particles/µl) or AAV2/PHP.eB-*hGfaABC*_*1*_*D*-DIO-TDP43-ΔNLS (1.3×10^13^ particles/µl). After 3–5 days, primary neurons were isolated and seeded on top of the transduced astrocytes as described above. Neural activity was recorded for 15–30 min daily between neuronal DIV 8–18 using the Maestro Pro MEA System (Axion BioSystems) and firing rates were analyzed using Neural Metric Software (Axion BioSystems). Some wells received the CXCR3 antagonist SCH 546738 (MedChemExpress #HY-10017; 12 nM).

### Slice electrophysiology

*Slice preparation:* Mice were deeply anesthetized with 5% isofluorane before being cardially perfused with ice-cold and oxygenated (95% O_2_/5% CO_2_) sucrose cutting solution. Sucrose cutting solution contained (in mM): 87 NaCl, 75 sucrose, 2.5 KCl, 1.25 NaH_2_PO_4_, 0.5 CaCl_2_, 25 NaHCO_3_, 1.3 ascorbic acid, and 10 D-glucose. The mice were quickly decapitated and the brain was extracted in ice-cold sucrose cutting solution. Coronal slices (350 μm thick) were made on a vibrating blade microtome (Leica VT1200s) while submerged in ice-cold and oxygenated sucrose cutting buffer. Slices were transferred to a heated (∼35^°^C) incubation chamber containing artificial cerebral spinal fluid (ACSF), which consisted of (in mM): 124 NaCl, 2.5 KCl, 1.5 MgSO_4_, 1.25 NaH_2_PO_4_, 2.5 CaCl_2_ and 26 NaHCO_3_. After approximately 30 minutes, the incubation chamber was allowed to equilibrate to room temperature for at least an additional 30 minutes.

#### Field potential recordings

For recordings, slices were transferred to a stage mounted holding chamber on an upright BX51W1 microscope (Olympus). The chamber was superfused (2–3 ml/min) with oxygenated (95% O_2_/5% CO_2_) and heated (∼35^°^C) ACSF. Recordings were obtained using a Multiclamp 200B amplifier (Molecular Devices) and filtered at 2 kHz, digitized at 10 kHz, and acquired with Clampex 10.7 (Molecular Devices). Micropipettes were made with borosilicate glass pulled to a resistance of 3.5 – 5.5 MΩ on a Flaming/Brown P-1000 micropipette puller (Sutter Instruments) and filled with the same ACSF. These were placed in the stratum radiatum of CA1 and a concentric bipolar stimulating electrode (FHC) was placed within the same layer, upstream of the Schaffer collaterals. Care was taken to keep the recording pipette and stimulating electrode as far apart as possible (at least 200 μm) to help isolate the stimulus artifact, the fiber volley, and the field potential. For stimulus intensity/field potential slope relationship, once a minimal current to reliably evoke a fiber volley and field potential was established, stimulus current was systematically increased and the subsequent fiber volley and field potential was recorded. For paired-pulse ratio recordings, stimulus intensity was set to approximately half-maximal intensity. Recordings were analyzed using custom code in Matlab (MathWorks), Excel (Microsoft) or Prism (GraphPad).

### VSV production, purification, and quantification

293T cells (ATCC, #CRL-3216) were plated at 1×10^6^ cells per well in 6-well plates. The following day, the cells were rinsed with serum-free medium and transfected with a mixture of plasmids encoding the rVSV antigenome, rVSV-ΔG-Luciferase (500 ng, Kerafast, #EH1007), and the rescue plasmids pCAG-VSVP (Addgene, Plasmid #64088), pCAG-VSVN (Addgene, Plasmid #64087), pCAG-VSVM (Addgene, Plasmid #64086), pCAG-VSVL (Addgene, Plasmid #64085), pCAG-VSVG (Addgene, Plasmid #64084) and pCAG-T7pol (Addgene, #59926). Lipofectamine 3000 (Thermo, #L3000001) was used for transfection according to the manufacturer’s instructions. After 48 h, the supernatant was collected, filtered through a 0.4-μm filter, and used to infect VSV-G–expressing cells for amplification. To amplify rescued rVSV-ΔG-Luciferase, 5 × 10^6^ 293T cells were plated per 10-cm dish in 10 ml of growth medium or 1.2 ×10^7^ 293T cells were plated per 15-cm dish. The cells were transfected with 5 μg (in 10-cm dish) or 12.5 μg (in 15-cm dish) of pCMV-VSV-G expression plasmid (Addgene, Plasmid #8454) using Lipofectamine 3000. The following day, the transfected cells were infected with the rescued virus, and 24–48 h later the supernatant was collected, centrifuged at 350 x g to clarify, and filtered through a 0.22-μm filter. VSV stock titer was quantified by serial dilution followed by infection. Briefly, 293T cells were plated at 2 × 10^5^ cells per well in 24-well plates. The following day, cells were rinsed with serum-free medium and infected with serially diluted VSV. Media was changed 2 h post-infection and the cells were fixed and immunostained 48 h post-infection to detect firefly luciferase (Abcam #ab181640, RRID # AB_2889835).

### VSV and adenovirus infections

Primary astrocytes from NTG and doubly transgenic ΔNLS mice were transfected with 0.66 µg of low molecular weight poly(I:C) (InvivoGen # tlrl-picw) per ml culture medium at DIV 8 using Lipofectamine 3000 5 h before infection with vesicular stomatitis virus (VSV) at 100 MOI or with adenovirus-eGFP (Ad5CMV-eGFP, lot #ad3586, Viral Vector Core Facility, Carver College of Medicine, University of Iowa) at indicated MOIs (see figures). A change of media was made 2 h after viral infections and the cells were collected 24 h after infection. VSV levels were measured by RT-qPCR. eGFP fluorescence was analyzed after cultures were fixed with 4% PFA in PBS and stained with DAPI. Fluorescence intensity (integrated density), area, and percent of total area were extracted using FIJI. Data are represented as eGFP normalized to the total DAPI-positive area and percentage of DAPI-positive area per condition.

### HSV-1 infections

HSV-1 H129-eGFP strain was generously provided by Dr. Lynn Enquist (Princeton University, Princeton, NJ). Viral stocks were grown on Vero E6 cells (ATCC, CRL-1586) maintained in Dulbecco’s minimum essential media (DMEM) with 10% fetal bovine serum (FBS). Standard plaque assays were performed to titer HSV-1 H129-eGFP stocks and quantify viral load in conditioned medium from infected primary astrocytes. Briefly, stocks or media samples were serially diluted in DMEM supplemented with 2% FBS and used to infect Vero cells. After 3 h, cells were washed three times with PBS and further incubated in DMEM with 2% FBS, antibiotics, and 1.5% methylcellulose (37°C). After 48 h, cultures were fixed with 4% PFA in PBS and counterstained with Hoechst 33342 (Thermo). Fluorescent signal was used to detect plaques of infected cells and quantify viral titers. Cell culture plates were imaged on a BX-X710 microscope (Keyence) with a 20X objective (Nikon). Images were stitched with BZ-X Analyzer Software (Keyence) and used to count plaques.

### Immunocytochemistry

All immunostaining steps were performed at ambient temperature unless specified otherwise. Briefly, cells were fixed with 4% paraformaldehyde and 4% sucrose in PBS for 10 min, rinsed four times with PBS (Corning) with 0.01% Triton X-100, and blocked and permeabilized in 5% normal goat serum (Jackson ImmunoResearch) or 5% normal donkey serum (Jackson ImmunoResearch) in 0.2%–0.3% Triton X-100 in PBS for 1 h. Cells were incubated overnight at 4°C with the following primary antibodies diluted in 1% BSA, 2% normal donkey serum, or 2% normal goat serum in 0.2%–0.3% Triton X-100 in PBS: mouse anti-PSD-95 (1:1000; Antibodies Incorporated # 75-028; RRID #AB_2292909), guinea pig anti-synaptophysin 1 (1:750; Synaptic Systems #101 004; RRID #AB_1210382), rabbit anti-bassoon (1:1000; Synaptic System #141 003 RRID #AB_887697), rabbit anti-RFP (1:500; Abcam #ab34771; RRID # AB_777699), human-specific mouse anti-TDP-43 (1:500; clone 6H6E12; ProteinTech # 60019-2-Ig; RRID #AB_2200520), or goat anti-GFAP (1:500; Abcam #ab53554; RRID #AB_880202). Cells were rinsed four times with PBS with 0.01% Triton X-100 and incubated for 1 h with the following AlexaFluor-conjugated secondary antibodies diluted in 1% BSA, 2% normal donkey serum, or 2% normal goat serum, and 0.2%–0.3% Triton X-100 in PBS (1:500; ThermoFisher #A11073, A31571, A31572; RRID # AB_2534117, AB_162542, AB_162543). Cells were rinsed twice with PBS with 0.01% Triton X-100 and twice with PBS before imaging.

### Immunohistochemistry

Postmortem human brain tissue blocks or sections from non-demented controls, AD, or bvFTD cases were obtained from the Neurodegenerative Disease Brain Bank at UCSF (San Francisco, CA) and the Banner Sun Health Research Institute Brain and Body Donation Program of Sun City, Arizona. Case details are listed in Supplementary Table 1. Formalin-fixed tissue blocks were rinsed in PBS and incubated in 30% sucrose for 3–5 days at 4°C before sectioning.

Mice were anesthetized with Avertin (2,2,2-tribromoethanol, 400–600 mg/kg body weight, Acros Organics) and transcardially perfused for 2.5 min with 0.9% saline before hemibrains were removed and stored in fixative (4% paraformaldehyde in PBS) overnight at 4°C on a rocking platform. Hemibrains were subsequently incubated in cryoprotectant (30% sucrose in PBS) for at least 48 h before sectioning.

Human and mouse brain tissue was sectioned (30 μm-thick sections) using a SM2010 R sliding microtome (Leica) equipped with a BFS-3MP freezing stage and cooling unit (Physitemp, Clifton, NJ). Free-floating sections were collected into cryopreservative (30% ethylene glycol, 30% glycerol in PBS) for long-term storage at -20°C.

Human tissue was immunolabeled by rinsing sections in PBS and permeabilizing overnight in PBS containing 0.5% Triton X-100 (PBS-T). Antigen retrieval was performed for 15 min in hot 0.1 M citrate buffer at pH 6.0, followed by incubation for 15 min with 3% hydrogen peroxide and 10% methanol in PBS. Sections were blocked for 2 h in 10% normal donkey serum (Jackson ImmunoResearch) and 2% non-fat dry milk and incubated overnight on a rocking platform with primary antibodies in 3% normal donkey serum. To minimize autofluorescence, sections were incubated for 20 minutes with 0.2 µm-filtered 0.3% Sudan Black B (Acros) in 70% ethanol and then incubated with secondary antibodies in 3% normal donkey serum for 2 h. The following primary antibodies were used for human tissue: pan-specific rabbit anti-TDP-43 (1:500; ProteinTech #10782-2-AP; RRID #AB_615042), mouse anti-GFAP (1:500; Millipore; #MAB3402B; RRID #AB_10917109), mouse anti-Aldh1L1; Clone N103/39 (1:1000; Millipore; #MABN495; RRID #AB_2687399), and AlexaFluor-conjugated secondary antibodies (1:250; ThermoFisher; listed below).

Double or triple immunolabeling of free-floating mouse sections was performed with minor modifications depending on the antibodies used. All steps were performed at ambient temperature unless specified. Cryopreserved sections were rinsed in PBS, permeabilized for 30 min or longer in PBS containing 0.5% Triton X-100 (PBS-T), blocked with 10% Normal Donkey or Goat serum (Jackson ImmunoResearch) in PBS-T for 1–2 h, incubated in primary antibodies in 3% serum in PBS-T for up to 48 h at 4°C, and rinsed with PBS-T. Sections were protected from light in all subsequent steps. Tissue was incubated with fluorescent secondary antibodies in 3% serum in PBS-T for 2 h, rinsed with PBS-T, mounted, and dried on Superfrost glass slides (VWR #75799-266) before sealing #1.5 coverglass (VWR #89239-734) with Vectashield antifade media containing DAPI (VWR #101098-050). When necessary, Prolong Diamond Antifade Mounting Media (ThermoFisher #P36970) replaced Vectashield mounting media to minimize quenching of AlexaFluor 647-conjugated antibodies. Slides were allowed to set overnight before acquiring images.

Unless co-labeled with an antibody requiring modified protocols described below, the following primary antibodies were used according to the general protocol: pan-specific rabbit anti-TDP43 (1:1000; ProteinTech #10782-2-AP; RRID #AB_615042), goat anti-CXCL10 (1:150; R&D Systems #AF-466-NA; RRID # AB_2292487), rabbit anti-GFAP (1:1000; Millipore-Sigma #G9269; RRID #AB_477035), mouse biotin-anti-GFAP (1:1000; Millipore-Sigma #MAB3402B; RRID #AB_10917109), rabbit anti-NeuN (1:1000; Millipore-Sigma #ABN78; RRID #AB_10807945), mouse anti-NeuN (1:1000; Millipore-Sigma #MAB377; RRID # AB_2298772), rabbit anti-Iba1 (1:1000; Wako #019-19741; RRID #AB_839504) and guinea pig anti-synaptophysin-1 (1:750; Synaptic Systems #101 004; RRID #AB_1210382). Following the permeabilization step, some antibodies required an antigen-retrieval step of 15 min in hot citrate buffer as described previously (*42*). These antibodies were mouse anti-viperin/*Cig5* (1:50; Abcam # ab107359, RRID #AB_10888107), mouse anti-synaptotagmin-2 (1:200; Developmental Studies Hybridoma Bank #znp-1-c), rabbit anti-CXCL9 (1:50; Abcam #ab202961), rabbit anti-CXCR3 (1:200; ProteinTech #26756-1-AP), rabbit anti-glutamine synthetase (1:250; ThermoFisher #701989, RRID #AB_2633045) and goat anti-PSD-95 (1:500; Abcam #ab12093; RRID # AB_298846).

For labeling with certain mouse monoclonal antibodies, the serum in the blocking and antibody steps was replaced with reagents from the Mouse-on-Mouse (M.O.M.) Basic Immunodetection Kit following vendor instructions (Vector Labs #BMK-2202). These antibodies were mouse anti-viperin/*Cig5*, mouse anti-synaptotagmin-2, human-specific mouse anti-TDP-43 (1:7,000; clone 6H6E12; ProteinTech # 60019-2-Ig; RRID #AB_2200520) and mouse anti-gephyrin (1:200; Synaptic Systems #147 011; RRID #AB_887717). Note that antigen retrieval with hot citrate buffer eliminated labeling by the human-specific mouse anti-TDP-43 antibody. AlexaFluor-conjugated secondary antibodies were obtained from ThermoFisher (1:500; #A11055, A21202, A21206, A21432, A21435, A31570, A31571, A31572; RRID #AB_2534102, AB_141607, AB_2535792, AB_141788, AB_2535856, AB_2536180, AB_162542, AB_162543).

### Microscopy and image analyses

Primary neurons were evaluated for synaptic content using a 40X objective on an ImageXpress MICRO Confocal Automated High-Content Analysis System (Molecular Devices) at the Weill Cornell Medicine Automated Optical Microscopy Core Facility. Briefly, four regions of interest (ROIs) were imaged per well and images were processed to assess the number and total area of synaptophysin-1 or PSD-95-positive puncta using FIJI (*139*). Briefly, images of individual channels were background subtracted, stringently thresholded to the brightest 5–10% of all pixels, and particles equal or larger than 1 µm in length were counted. The total area and number of thresholded puncta were measured for each image.

To measure astrocytic TDP-43 in human brain tissue, postmortem hippocampal tissue was immunostained with anti-TDP-43 and anti-GFAP antibodies and imaged on a BX-X710 microscope (Keyence) with a 40X objective (Nikon) and astrocytes were analyzed within the dentate gyrus and Cornu Ammonis regions. Supplemental Tables 1–2 provide detailed information about the human cases and the numbers of cells analyzed per case. Image analysis was performed using FIJI (*139*). Single-channel immunostaining did not reveal bleed-through to other channels (data not shown). In addition, the patterns and intensities for TDP-43 and GFAP were not consistently colocalized across cases and individual cells.

Astrocytes were defined by drawing ROIs encompassing primary processes and cell soma with strong GFAP fluorescence together with overlapping or adjacent DAPI-positive nucleus. ROIs were extracted and the intensity of nuclear TDP-43 immunoreactivity was measured for pixels within a DAPI-thresholded mask. The intensity of extra-nuclear TDP-43 immunoreactivity in GFAP-positive ROIs was measured for pixels in a GFAP-thresholded mask after eliminating the nuclear area defined by the DAPI mask. Intensities were corrected for background fluorescence in each image by subtracting the mean intensity of five circular ROIs drawn in areas that did not have GFAP, TDP-43, or DAPI labeling above background levels.

To evaluate cell-specific localization and intensity of protein expression in mouse brain sections, slides were imaged on a BX-X710 microscope (Keyence) with a 20X objective (Nikon) using the tiling function. Images were stitched with BZ-X Analyzer Software (Keyence) and further processed for brightness and contrast using FIJI. To further evaluate cell localization of proteins in mouse brain sections, z-stacks of immunostained tissue were acquired on a Zeiss LSM 880 Laser Scanning Confocal Microscope with a 63X objective and three-dimensional renderings of maximal projections were made using Imaris software (Oxford Instruments).

Localization of CXCR3 protein in neuronal cell bodies and synaptic compartments was evaluated using a Zeiss LSM 880 Laser Scanning Confocal Microscope. Images of the CA1 stratum radiatum region of the hippocampal formation were acquired using a 63X objective, 4X zoom, 16-line averaging, and a pixel dwell time of 5.3 µs. Filter and detector configurations were optimized using single-antibody controls and each of the three channels was imaged sequentially to further minimize potential cross-bleed. Images were analyzed using FIJI to measure the number, area, and intensity of puncta that were immunoreactive for CXCR3 and NeuN, synaptophysin-1, synaptotagmin-2, PSD-95, or gephyrin.

Briefly, images of individual channels were background subtracted and noise was removed using the “despeckle” function. Images were stringently thresholded to the brightest 5– 10% of pixels and particles equal or larger than 1 µm in length were counted. The total area, mean size, and mean intensity of thresholded puncta were measured for each image. The fractional overlap of CXCR3 and each synaptic marker was measured from processed images as Mander’s coefficient using the Just Another Colocalization Plugin (JACoP) (*140*). Total of 9–12 images from 3–4 mice were analyzed per genotype.

### Western blotting

For cell cultures, wells were rinsed twice with ice-cold PBS before aspirating buffer and lysing directly with ice-cold 1X RIPA Buffer (Thermo Fisher #89900) containing 1X cOmplete Protease Inhibitor Cocktail (Millipore Sigma #11836153001) and 1% each of Phosphatase Inhibitor Cocktails 2 and 3 (Millipore Sigma #P5726 and #P0044). Cells were scraped, collected in 1.5 ml Eppendorf tubes, sonicated on ice for 5 s at 10% power with a probe sonifier (Branson), centrifuged for 5 min at 10,000 g at 4°C, and assayed for protein content using a detergent-compatible Bradford assay (Thermo Fisher #23246).

For brain tissue, approximately 30 mg of mouse hippocampal tissue was dissected from flash-frozen forebrain in ice-cold PBS under a dissecting scope (AmScope) and re-frozen on dry ice in 1.5 ml Eppendorf tubes. Frozen samples were thawed on ice for 1–2 min before adding 150 µl ice-cold lysis buffer to each tube (RIPA with protease and phosphatase inhibitors). Samples were immediately homogenized with a Fisherbrand Bead Mill 24 (Fisher Scientific #15-340-163) for 40 s at a speed setting of 5 in a pre-chilled adaptor tube rack. Samples were centrifuged for 2 min at 1000 g at 4°C before sonication in an ice-chilled EpiSonic 2000 water bath (5 s on, 2 s off, 5 min total, amplitude 40). RIPA-soluble extracts were clarified by centrifugation at 100,000 g for 30 min at 4°C in a Beckman Ultracentrifuge and protein content was measured using a detergent-compatible Bradford assay (Thermo Fisher).

For tissue and cell culture extracts, 30 µg or 20 µg RIPA-soluble lysates, respectively, were resolved on Bis-Tris SDS-PAGE gels (ThermoFisher) and transferred onto nitrocellulose membranes using an iBlot2 Western blotting system or a Mini Blot Module (ThermoFisher). Membranes were blocked with 5% non-fat milk or 5% bovine serum albumin (BSA; VWR #97062-904) in TBS before probing overnight at 4°C with primary antibodies diluted in TBS containing 0.2% Tween-20 (TBS-Tw). Primary antibodies were raised against TDP-43 (1:2000 pan-specific rabbit anti-TDP-43 ProteinTech #10782-2-AP; RRID # AB_615042 or 1:2000 human-specific mouse anti-TDP-43 ProteinTech #60019-2-Ig; RRID #AB_2200520), β-actin (1:2000; rabbit; Sigma #A2066; RRID #AB_476693), γ-tubulin (1:1,250; mouse; Sigma #T5326; RRID #AB_532292); NF-KB (1:1000 mouse anti-total NF-κB; Cell Signaling Technology #6956; RRID #AB_10828935 or 1:1000 rabbit anti-phospho NF-κB-S536; Cell Signaling Technology #3033; RRID #AB_331284); Akt (1:250 mouse anti-total Akt; Cell Signaling Technology #2920; RRID #AB_1147620 or 1:2500 rabbit anti-phospho Akt (S473); Abcam ab81283; RRID # AB_2224551); ERK1/2 (1:1000 mouse anti-total ERK1/2; Cell Signaling Technology #4696; RRID # AB_390780 or rabbit anti-phospho ERK1/2 (T202, Y204); Cell Signaling Technology #9101; RRID #AB_331646). STAT3 (1:1000 mouse anti-total STAT3; Cell Signaling 9139S RRID # AB_ 331757 or 1:1000 rabbit anti-phospho STAT3 (Y705); Cell Signaling 9145S RRID #AB_ 2491009).

After overnight incubation in primary antibodies, all blots were rinsed with TBS-Tw and probed for 1 h with IR Dye 680RD donkey anti-mouse (1:15,000; LI-COR #926-68072, RRID #AB_2814912) and IR Dye 800CW donkey anti-rabbit (1:15,000; LI-COR #926-32213; RRID #AB_621848) in TBS-Tw with 3% BSA. Blots were rinsed twice with TBS-Tw, once with TBS, and dried for at least 20 min before scanning on the Odyssey CLx imaging system (LI-COR). Expression levels were quantified using LI-COR Image Studio software.

### Microfluidic qPCR

RNA was extracted using the RNeasy Mini Kit with on-column DNase treatment following manufacturer instructions (Qiagen #74106, #79256). Cultured primary cells were rinsed once with ice-cold PBS, scraped in freshly prepared extraction buffer and frozen at -80°C until extracted. Saline-perfused, microdissected mouse brain tissue was frozen on dry ice and stored at -80°C until RNA extraction. Tissue was homogenized in fresh extraction buffer using a bead mill for 20 s at a speed setting of 5 in a pre-chilled adaptor tube rack.

The number of VSV viral copies was determined by RT-qPCR using PowerUp SYBR Green Master Mix (Thermo Fisher #A25741) according to manufacturer’s instructions. RT-qPCR was performed in a CFX96 Touch Real-Time PCR Detection System (BioRad). All primer sequences are detailed in Supplemental Table 3.

Microfluidic RT-qPCR was performed similarly to described protocols (*141-143*). Briefly, cDNA was synthesized with Protoscript First Strand Synthesis Kit (New England Biolabs #E6300L) and pre-amplified for 14 cycles against a pool of primers (Supplemental Table 3) using PreAmp Grandmaster mix (TATAA Biocenter, Sweden #TA05) before exonuclease I treatment (New England Biolabs #M0293L). Pre-amplified cDNA was diluted at least 5-fold with nuclease-free water and mixed with SsoFast EvaGreen with Low ROX (BioRad #1725211) and chip-specific DNA Sample Reagents before loading into primed Flex Six or 96.96 Dynamic Array chips (Fluidigm #100-6308, BMK-M-96.96). Individual primers were mixed with DNA assay reagent (Fluidigm) and loaded into chip inlets. Chips were primed and loaded using an IFC Controller HX (Fluidigm) before measuring and analyzing amplification and melting curves on a BioMark HD System (Fluidigm). Cycle of quantification (Cq) values were thresholded equally for all inlets across each chip run and normalized to the average of reference genes (*Actb* and *Gapdh* for tissue samples; *Actb* and/or *Tbp* for cultured cells) before determining ddCq and fold-change relative to experimental control groups.

### Statistical analyses

Statistical specifications are reported in the figures and corresponding figure legends. Data are presented as mean ± S.E.M. All statistical tests were performed using GraphPad Prism 8, except Fisher’s exact test, which was performed using IBM SPSS Statistics for Windows, Version 24.0. The criterion for data point exclusion was established during the design of the study and was set to values above or below two standard deviations from the group mean. Two-sided Student’s *t* test was used to determine statistical significance between two groups. Welch’s correction was used to account for unequal variances. Differences among multiple groups were assessed by one-way or two-way ANOVA or mixed-effects model followed by Dunnett’s or Bonferroni’s multiple comparisons post-hoc tests, as specified in the legends. Null hypotheses were rejected at p < 0.05.

## Acknowledgements

We thank S. Tymchuk, C. Heisenberg, S. Shatela, and T. Chowdhury for technical support, and M. Garvey, G. Coronas-Samano, L.J. Metakis, and E. Spencer for administrative support. We are grateful to the Banner Sun Health Research Institute Brain and Body Donation Program of Sun City, Arizona, and the Neurodegenerative Disease Brain Bank at the University of California, San Francisco for the provision of human tissue samples. The Banner Brain and Body Donation Program received support from the National Institute of Neurological Disorders and Stroke, U24 NS072026 National Brain and Tissue Resource for Parkinson’s Disease and Related Disorders; National Institute on Aging, P30 AG19610 Arizona Alzheimer’s Disease Core Center; Arizona Department of Health Services, Arizona Alzheimer’s Consortium; Arizona Biomedical Research Commission, Arizona Parkinson’s Disease Consortium; and the Michael J. Fox Foundation for Parkinson’s Research. The Neurodegenerative Disease Brain Bank at the University of California, San Francisco received support from NIH grants P01AG019724 and P50AG023501, the Consortium for Frontotemporal Dementia Research, and the Tau Consortium. All schematics were generated using BioRender.com.

## Funding

NIAID grant 2R01AI107301 (RES)

NIDDK grant R01DK121072 (RES)

NCI grant R01CA234614 (RES)

Irma T. Hirschl Trust Research Award Scholar (RES)

NIDCD grant DC012557 (RCF)

NIH BRAIN Initiative grant NS107616 (RCF)

Howard Hughes Medical Institute Faculty Scholarship (RCF)

NINDS/NIA grant 1R01NS118569 (AGO)

NIA grant R00AG048222 (AGO)

Alzheimer’s Association (AGO)

Leon Levy Foundation (AGO)

## Author contributions

ALM, ALO, and AGO designed the study and drafted the manuscript; ALM, SM, FP, CZ, SJ, ALO, and AGO performed experiments and contributed to the acquisition and analysis of data; YB and RES performed or assisted with viral pathogen experiments; SCS and RCF performed or assisted with slice electrophysiology.

## Data and materials availability

Raw data are available from the corresponding authors upon request. All materials are available upon request or commercially.

