## Supplemental Figures and Tables for "Astrocytic TDP-43 dysregulation impairs memory by modulating antiviral pathways and interferon-inducible chemokines"

**This file includes:**

Figs. S1 to S8

Tables S1 to S3

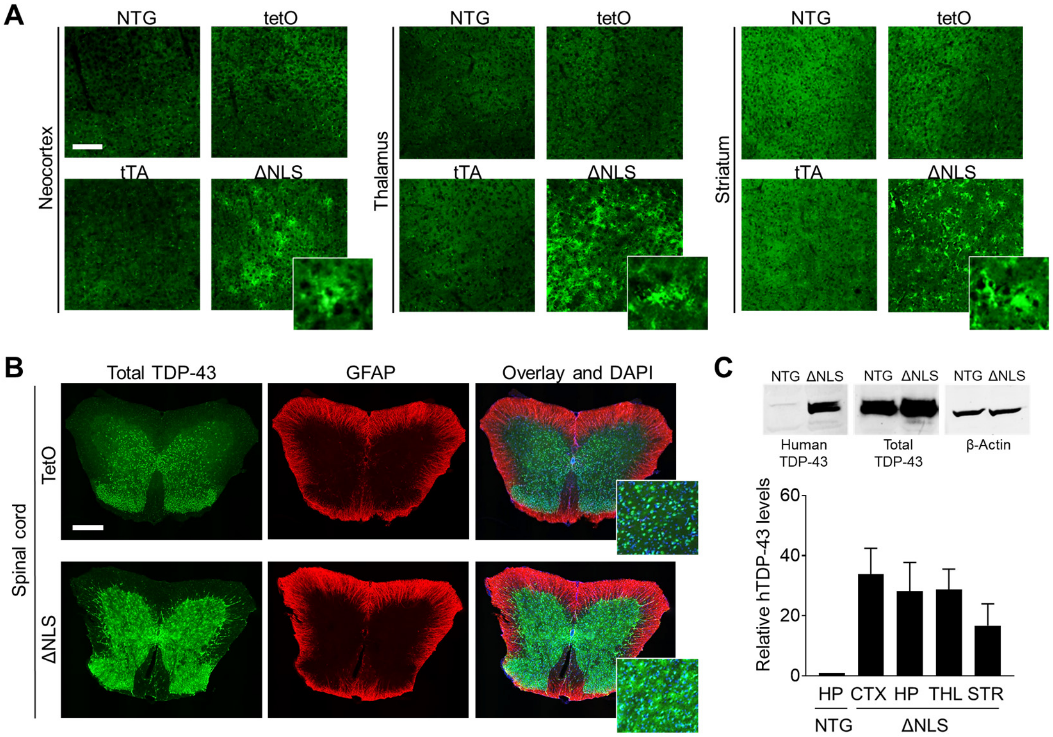


**Fig. S1. Additional characterization of transgenic mice with inducible TDP-43 alterations in astrocytes.**

(**A**) Representative images of human-specific TDP-43 (green) immunoreactivity in indicated brain regions of 3-month-old nontransgenic (NTG) controls, singly transgenic *hGFAP*-tTA (tTA) and *tetO*-TDP43-ΔNLS (TetO) controls, and doubly transgenic ΔNLS mice. (**B**) Immunolabeling for total (mouse and human) TDP-43 levels (green) and the astrocyte marker GFAP (red) in the spinal cord of TetO controls and doubly transgenic ΔNLS mice. DAPI (blue) was used to visualize nuclei. Insets in (A and B) show magnified views. Scale bars: 100 µm (A), 300 µm (B). (**C**) Western blot images (top) and quantification (bottom) for human (h) and total (t) TDP-43 in different brain regions of NTG controls and ΔNLS mice. Representative images show Western blotting of hippocampal samples for indicated proteins. TDP-43 levels were normalized to β-actin levels. n = 4–5 mice per brain region.

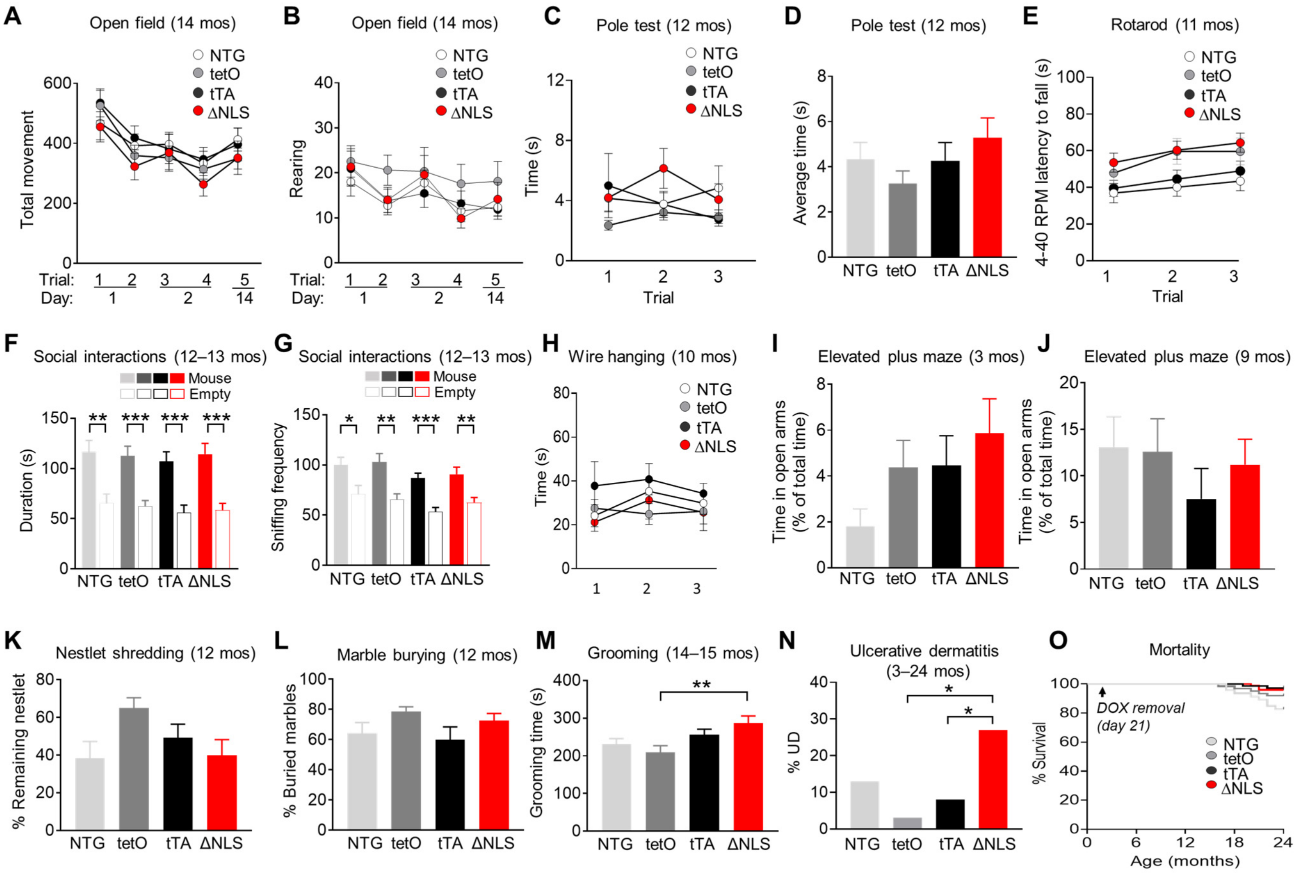


**Fig. S2. Additional behavioral characterization of transgenic mice with inducible TDP-43 alterations in astrocytes.**

(**A**–**N**) Nontransgenic (NTG) controls, singly transgenic *hGFAP*-tTA (tTA) and *tetO*-TDP43-ΔNLS (TetO) controls, and doubly transgenic ΔNLS mice were assessed for exploratory behavior in the open field (**A** and **B**), pole test (**C** and **D**), motor behavior on the Rotarod (**E**), interactions with a novel mouse as compared to an empty wire cup (Empty) as a measure of sociability (**F**–**G**), wire hanging test (**H**), anxiety-related behavior in the elevated plus maze (**I** and **J**), nestlet shredding (**K**), and marble burying in bedding (**L**). Mice were also assessed for time spent grooming after a water mist (**M**) and incidence of ulcerative dermatitis (**N**). Ages of mice (months, mos) are indicated in each panel. (**O**) Early mortality was assessed for up to 24 months of age. (A–O) Student's *t* test with Welch's correction: *$p < 0.05$, **$p < 0.01$, ***$p < 0.001$ vs. Empty (F–G); One-way ANOVA: $F(3, 64) = 3.63$, $p = 0.0175$ (M); Dunnett's post-hoc test: **$p < 0.01$ vs. tetO (M). Fisher's exact test (two-sided): $p = 0.023$ (overall); *$p = 0.035$ vs. tTA, $p = 0.007$ vs. tetO, $p = 0.16$ vs. NTG (N). n = 15–17, 33 females and 30 males (A and B); 13–15, 33 females and 24 males (C and D); 13–15, 29 females and 26 males (E); 12–15, 30 females and 30 males (F and G); 12–15, 29 females and 26 males (H); 13–18, 28 females and 30 males (I); 13–18, 28 females and 30 males (J); 12–15, 29 females and 26 males (K and L); n = 15–22, 37 females and 33 males (M); 35–40, 82 females and 68 males (N); 15–22**,** 37 females and 33 males (O) per genotype.

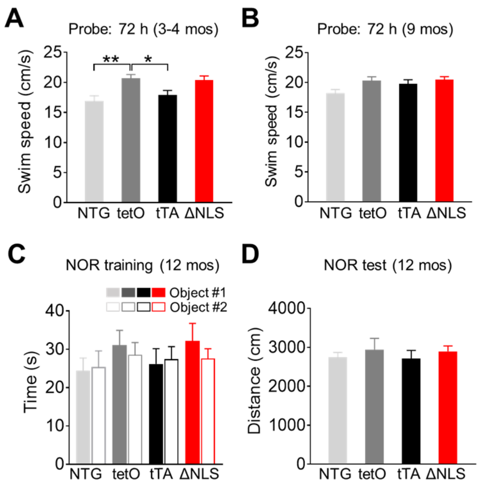


**Fig. S3. Morris water maze swim speeds and novel object recognition in transgenic mice with astrocytic TDP-43 alterations.**

(**A** and **B**) Nontransgenic (NTG) controls, singly transgenic *hGFAP*-tTA (tTA) and *tetO*-TDP43-ΔNLS (tetO) controls, and doubly transgenic ΔNLS mice were assessed for swim speeds in probe trials at indicated time-points (hours, h) after hidden platform training. (**C**) Exploration of objects during novel object recognition (NOR) training prior to the testing day. (**D**) Total exploration during NOR testing. Ages of mice (months, mos) are indicated in each panel. One-way ANOVA: $F(3, 48) = 6.1$, $p = 0.0013$ (A); $F(3, 66) = 2.98$, $p = 0.038$ (B); $F(3, 51) = 0.90$, $p = 0.448$ (C); $F(3, 4) = 0.335$, $p = 0.80$ (D). Dunnett's post-hoc test: *$p<0.05$, **$p<0.01$ vs. tetO. n = 12–15, 29 females and 26 males (A); 15–22, 37 females and 33 males (B); 12–15, 29 females and 26 males (C); 12–15, 29 females and 26 males (D) per genotype.

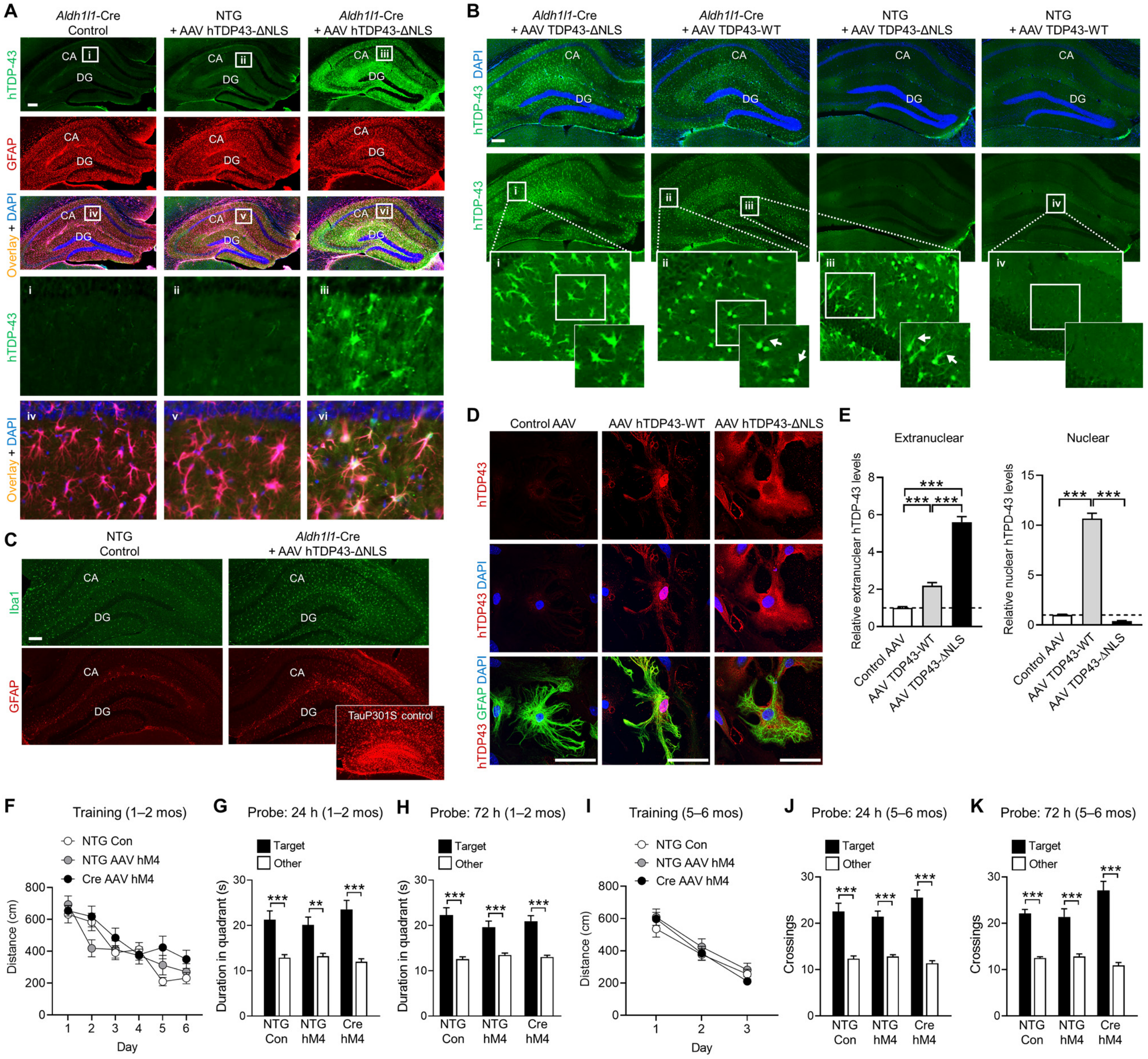


**Fig. S4. Additional characterization of AAV vectors in mice and cultured astrocytes.**

(**A**–**C**) Representative images of hippocampal immunolabeling for human TDP-43 (green), astrocytic marker GFAP (red) and/or microglial/macrophage marker Iba1 (green) in transgenic *Aldh1l1*-Cre or littermate nontransgenic (NTG) control mice injected with AAV PHP.eB-*hGfaABC*$_1$*D*-DIO-hTDP43-ΔNLS, AAV PHP.eB-*hGfaABC*$_1$*D*-DIO-hTDP43-WT, or saline (Control). Mice were assessed 9 months (A) or 3 weeks (B) after intracranial injections. Yellow indicates overlay of green and red channels. DAPI (blue) was used to visualize nuclei. Inset in (C) shows hippocampal GFAP immunoreactivity in a transgenic tau-P301S mouse as a positive control for hippocampal astrogliosis. Insets i–vi (A) and i–iv (B) show magnified views. (**D** and **E**) Representative images (D) and quantification (E) of immunolabeling for human TDP-43 in primary astrocytes derived from *Aldh1l1*-Cre mice and transduced with the indicated AAV vectors: control AAV PHP.eB (2.38E+10 Vg/well), AAV PHP.eB-*hGfaABC*$_1$*D*-DIO-hTDP43-WT (6.00E+10 Vg/well), or AAV PHP.eB-*hGfaABC*$_1$*D*-DIO-hTDP43-ΔNLS (2.90E+11 Vg/well). Levels of hTDP-43 protein expression were similar among AAV vectors as assessed by Western blotting (data not shown). Cultures were co-immunolabeled for human TDP-43 (red) and the astrocytic marker GFAP (green). Individual astrocytes were analyzed for levels of hTDP-43 immunoreactivity in different subcellular regions as indicated. One-way ANOVA: $F(2, 101) = 144.9$, p<0.0001 (extranuclear); $F(2, 101) = 424.2$, p<0.0001 (nuclear). Bonferroni's post-hoc test: ***p<0.001 vs. indicated group. n = 30–37 cells from 3 culture wells per condition. Scale bars: 200 µm (A–C), 50 µm (D). (**F**–**K**) Nontransgenic (NTG) or transgenic *Aldh1l1*-Cre (Cre) littermates were injected with AAV PHP.eB-*hGfaABC*$_1$*D*-DIO-hM4Di-mCherry (hM4) or saline (Con) at 5–6 months of age and tested in the Morris water maze at 1–2 or 5–6 months after AAV injections, as indicated. (**F** and **I**) Distance traveled to reach the platform during hidden platform training (four trials per session, one session per day). Repeated measures two-way ANOVA: $F(2, 50) = 1.96$, p = 0.15 for group effect (F); $F(2, 47) = 0.96$, p = 0.39 for group effect (I). n = 17 NTG/Con, 18 NTG/hM4, and 18 Cre/hM4 mice (27 females, 25 males) (F); n = 14 NTG/Con, 18 NTG/hM4, and 18 Cre/hM4 mice (25 females, 25 males) (I). (**G**–**H, J**–**K**) Probe trials were conducted 24 or 72 h after completion of training, as indicated. Durations in target and non-target (Other) quadrants. Student's *t* test with Welch's correction: **p < 0.01, ***p < 0.001 vs. Other.

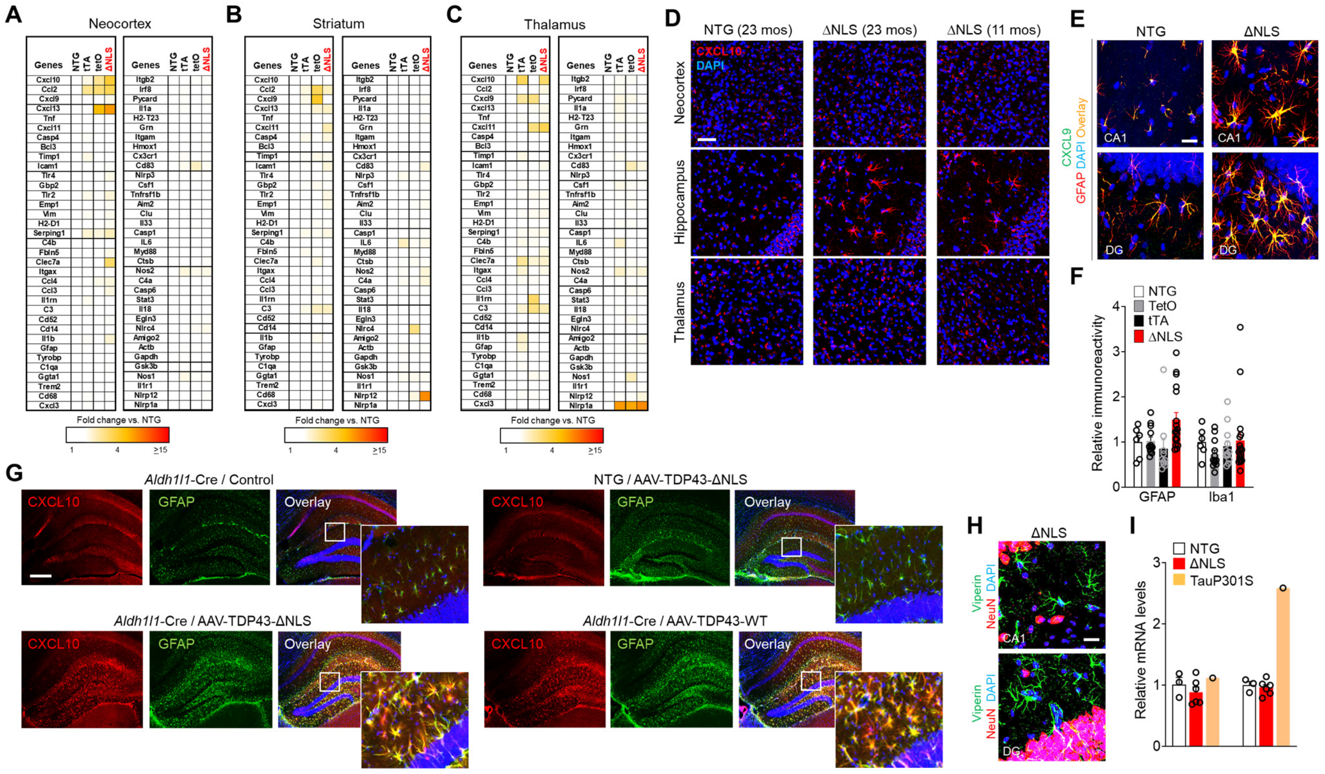


**Fig. S5. Additional characterization of gene expression in transgenic mice with astrocytic TDP-43 alterations.**

(**A**–**C**) Relative RNA levels in indicated brain regions of 11-month-old littermate nontransgenic controls (NTG), singly transgenic *hGFAP*-tTA (tTA) and *tetO*-TDP43-ΔNLS (TetO) controls, and doubly transgenic TDP-43-ΔNLS mice (ΔNLS). (**D**) Representative images of CXCL10 immunoreactivity (red) in indicated brain regions of NTG and ΔNLS mice at different ages. DAPI (blue) was used to visualize nuclei. (**E**) Representative images of CXCL9 immunoreactivity (green) in the CA1 region or dentate gyrus molecular layer (DG) of 11-month-old NTG controls and doubly transgenic ΔNLS mice. Sections were co-immunolabeled for the astrocyte marker GFAP. Yellow indicates overlay of green and red channels. DAPI (blue) was used to visualize nuclei. (**F**) Quantification of relative GFAP and Iba1 immunoreactivity (integrated brightness) in the hippocampal formation of 11-month-old NTG, TetO, tTA, and ΔNLS mice. One-way ANOVA: $F(3, 42) = 3.6$, $p = 0.021$ (GFAP); $F(3, 44) = 0.74$, $p = 0.53$ (Iba1). n = 6–18 regions of interest (ROIs) from 3–9 mice per genotype. (**G**) Representative images of CXCL10 (red) and GFAP (green) immunoreactivity in hippocampal sections from 13-month-old nontransgenic (NTG), transgenic *Aldh1l1*-Cre (Cre), or littermate nontransgenic (NTG) control mice injected with AAV PHP.eB-*hGfaABC$_1$D*-DIO-hTDP43-ΔNLS, AAV PHP.eB-*hGfaABC$_1$D*-DIO-hTDP43-WT, or saline (Control). Mice were assessed 9 months after intracranial injections. DAPI (blue) was used to visualize cell nuclei. Insets show magnified views of boxed regions. (**H**) Representative images of viperin immunoreactivity (green) in the CA1 region or dentate gyrus molecular layer (DG) of 11-month-old NTG controls and doubly transgenic ΔNLS mice. Sections were co-immunolabeled for the neuronal marker NeuN (red). Viperin did not overlap with NeuN. DAPI (blue) was used to visualize nuclei. (**I**) Hippocampal *Cgas* and *Sting1* RNA levels in 11-month-old NTG controls and ΔNLS mice, and transgenic TauP301S mouse as a positive control. n = 3–6 mice per genotype; n = 1 for TauP301S. Scale bars: 50 µm (D), 20 µm (E, H), 400 µm (G).

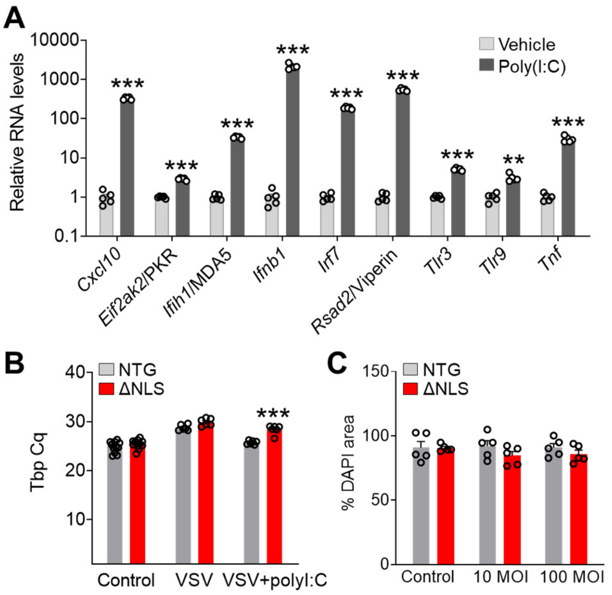


**Fig. S6. Additional characterization of antiviral responses in isolated astrocytes.**

(**A**) Primary astrocytes (DIV 9) derived from nontransgenic mice were acutely transfected with polyinosinic-polycytidylic acid (poly(I:C), 0.66 µg/µl in lipofectamine 3000) or treated with vehicle and were collected after 24 h for RNA analyses. RNA levels were normalized to TATA-binding protein (Tbp) and expressed as fold change relative to vehicle-treated controls. Student's *t* test with Welch's correction: **$p < 0.01$, ***$p < 0.001$ vs. Vehicle. n = 5 samples per treatment condition. (**B** and **C**) Primary astrocytes (DIV 9) derived from NTG or doubly transgenic ΔNLS mice were infected with vesicular stomatitis virus (VSV, 100 MOI) (B) or adenovirus tagged with eGFP at indicated MOIs (C). Some wells were also transfected with poly(I:C) as a control 5 h before viral infection. To compare cell density across treatments and genotypes, RNA levels of the control gene *Tbp* (B) or relative DAPI fluorescence (C) were assessed 24 h after infection. Two-way ANOVA: $F(2, 42) = 5.63$, $p = 0.007$ for interaction effect (B); $F(2, 24) = 0.54$, $p = 0.591$ for interaction effect (C). Bonferroni post-hoc test: ***$p < 0.001$ vs. NTG. n = 6–12 (B) and 5 (C) culture wells per genotype and treatment condition.

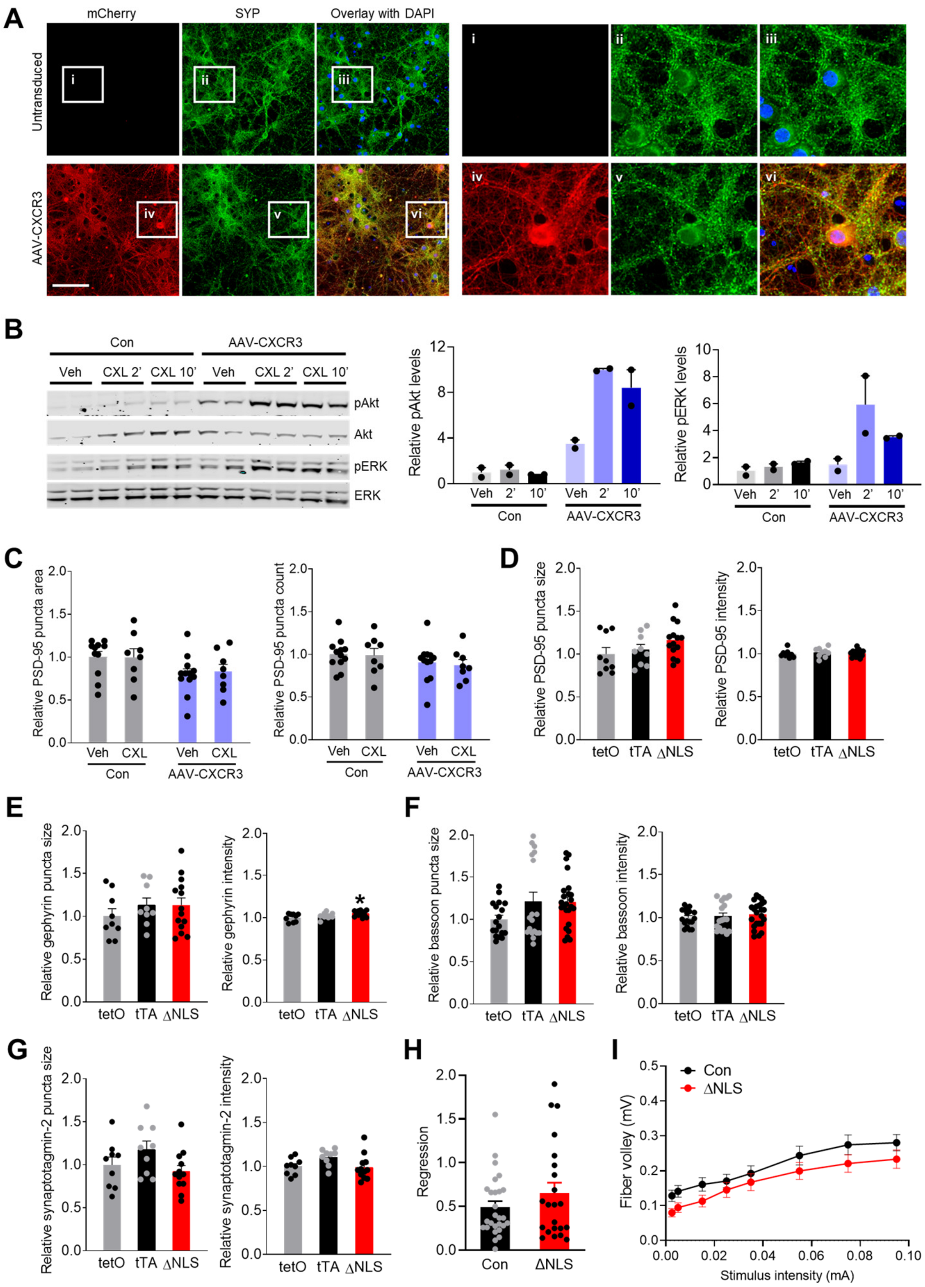


**Fig. S7. Characterization of *Cxcr3* transduction in isolated neurons and further analyses of synaptic markers and neuronal activity.**

(**A**) Representative images of mCherry (red) and synaptophysin (SYP, green) immunoreactivity in primary wild-type neurons 5 days after transduction with AAV DJ-*hSyn1*-mCherry-T2A-Cxcr3-2HA-neurexin1α (AAV Syn-CXCR3). DAPI (blue) was used to visualize cell nuclei. Yellow indicates overlap of green and red channels. Insets i–vi show magnified views. Scale bar: 100 µm. (**B**) Western blot images and quantification of phosphorylated and total Akt and ERK1/2 levels in Neuro2a cells untransfected (Con) or transfected with *hSyn1*-CXCR3 and then treated for 2 or 10 min with CXCL11 (CXL, 200 nM) or vehicle (Veh). n = 2 wells per treatment condition. (**C**) Quantification of PSD-95 immunoreactivity in primary wild-type neurons 5 days after transduction with AAV Syn-CXCR3 and treatment with CXCL11 (CLX, 200 nM) or vehicle (Veh) for 3 days. Control cultures (Con) were not transduced. Two-way ANOVA: $F(1, 35) = 0.055$, p = 0.816 for interaction effect (size); $F(1, 36) = 0.027$, p = 0.87 for interaction effect (count). n = 8–12 wells per condition. (**D**–**G**) Quantification of hippocampal immunoreactivity for synaptic markers PSD-95 (D), gephyrin (E), bassoon (F), and synaptotagmin-2 (G) in the CA1 region of singly transgenic controls (tetO and tTA mice) and ΔNLS mice. One-way ANOVA: $F(2, 29) = 1.86$, p = 0.17 (size); $F(2, 29) = 0.23$, p = 0.78 (intensity) (D); $F(2, 29) = 0.69$, p = 0.52 (size); $F(2, 29) = 4.38$, p = 0.02 (intensity) (E); $F(2, 57) = 2.33$, p = 0.11 (size); $F(2, 57) = 0.38$, p = 0.68 (intensity) (F); $F(2, 27) = 2.37$, p = 0.11 (size); $F(2, 27) = 2.94$, p = 0.07 (intensity) (G); Dunnett's post-hoc test: *$p < 0.05$ vs. tetO. n = 9–24 per genotype (D–G). (**H** and **I**) Hippocampal slice recordings in control and ΔNLS mice. (H) Linear regression between fEPSP slopes and fiber volley amplitudes. (I) Fiber volley amplitudes measured at increasing stimulus intensities as indicated. Mixed-effects model: $F(1, 51) = 2.61$, p = 0.11 for genotype effect.

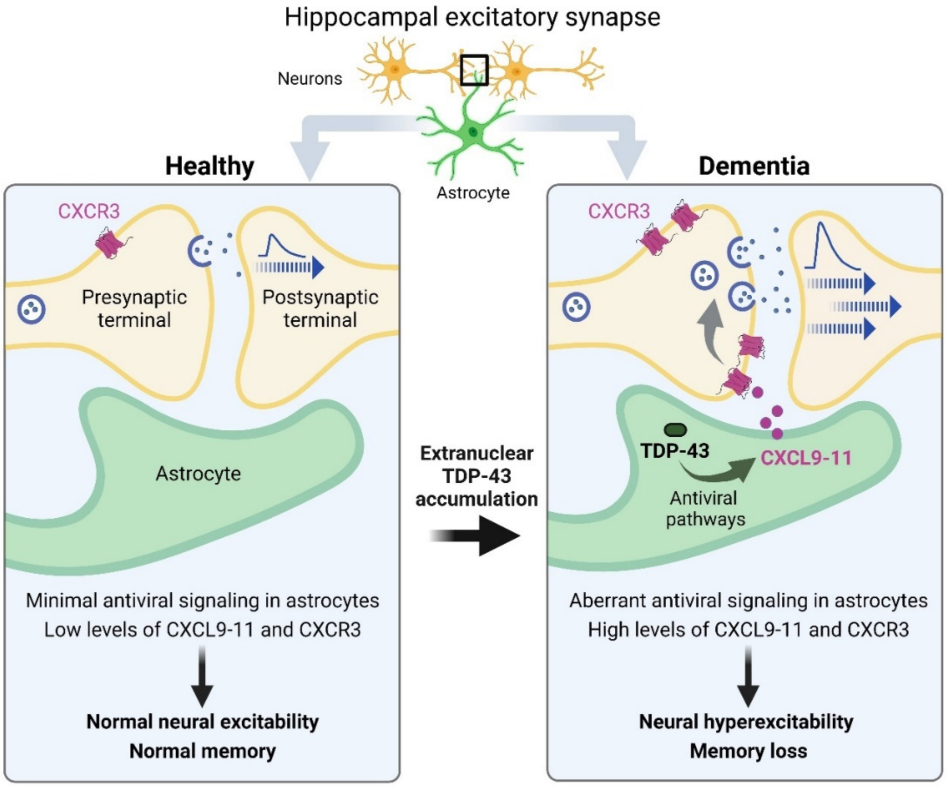


112 **Fig. S8. Summary of the main findings.**

**Table S1.** Clinicopathological information for human brain tissue used in Fig. 1.

| **Cases (n)** | **PMI (hours)** | **M/F Ratio** | **Age (years)** | **Individual NIA-Reagan Scores** |
|---|---|---|---|---|
| Con (9) | 4.8 +/- 2.2 | 2:1 | 80.2 +/- 7.3 | 4, 4, 4, 4, 1, 1, 1, 0, NA |
| AD (10) | 6.0 +/- 5.3 | 4:1 | 75.6 +/- 10.4 | 3, 3, 3, 3, 3, 3, 3, 3, 3, 3 |
| s-FTD (9) | 6.4 +/- 4.8 | 1:2 | 75.8 +/- 13.2 | 4, 4, 1, 1, 1, 0, NA, NA, NA |
| f-FTD (4) | 17.8 +/- 12.6 | 3:1 | 68.5 +/- 3.9 | NA, NA, NA, NA |

Values indicate mean +/- SD; Postmortem interval (PMI); Male-to-female ratio (M/F); Folstein Mini Mental State Examination (MMSE); NIA Reagan criteria for the postmortem diagnosis of Alzheimer's disease were developed jointly by the National Institute on Aging and the Reagan Institute (*1*). These criteria suggested that neuritic plaque density and Braak neurofibrillary staging offer probabilistic estimates of their likelihood for being responsible for dementia. Score 0 = not AD; 1 = low likelihood; 2 = intermediate likelihood; 3 = high likelihood; 4 = subject was not demented. NA: not available.

**Table S2.** Number of individual cells analyzed per case for human brain tissue used in Fig. 1.

| **Con (case #)** | **Cells (n)** | **s-AD (case #)** | **Cells (n)** | **s-FTD (case #)** | **Cells (n)** | **f-FTD (case #)** | **Cells (n)** |
|---|---|---|---|---|---|---|---|
| 1 | 41 | 1 | 26 | 1 | 21 | 1 (C9) | 24 |
| 2 | 24 | 2 | 16 | 2 | 28 | 2 (C9) | 21 |
| 3 | 36 | 3 | 18 | 3 | 27 | 3 (GRN) | 22 |
| 4 | 43 | 4 | 18 | 4 | 28 | 4 (GRN) | 18 |
| 5 | 16 | 5 | 12 | 5 | 28 | | |
| 6 | 18 | 6 | 73 | 6 | 68 | | |
| 7 | 18 | 7 | 20 | 7 | 16 | | |
| 8 | 23 | 8 | 24 | 8 | 24 | | |
| 9 | 17 | 9 | 18 | 9 | 21 | | |
| | | 10 | 24 | | | | |
| Total: | 236 | | 249 | | 261 | | 85 |

**Table S3.** Mouse gene primer sequences used for microfluidic qPCR.

| Gene | Forward Primer | Reverse Primer |
|---|---|---|
| Actb | CCCTAAGGCCAACCGTGAAA | AGCCTGGATGGCTACGTACA |
| Aim2 | TGGGCTGTTTAAAGTCCAGAA | CACCTCCATTGTCCCTGTT |
| Amigo2 | GCAGAGAGCTGCTGTGTC | GTGGGGCACATTCCTGAG |
| Bcl3 | GTGGAGAACAACAGCCTGAAC | CATCTGAGCGTTCACGTTGG |
| C1qa | ATGGGGCTCCAGGAAATCC | TCCCCTGGGTCTCCTTTAAAAC |
| C3 | ACCTTACCTCGGCAAGTTTCT | TTGTAGAGCTGCTGGTCAGG |
| C4a | CAGCACCTTTGTCAAGGTTACA | GACAAAGCTCCTTCAGAGCC |
| C4b | TCTCACAAACCCCTCGACAT | AGCATCCTGGAACACCTGAA |
| Casp1 | GGAGGACATCCTTCATCCTCA | GCAAAACTTGAGGGTCCCA |
| Casp4 | CCAGACATTCTTCAGTGTGGA | CTGGTTCCTCCATTTCCAGA |
| Casp6 | GCTCAAAATTCACGAGGTGTC | TCGTATGCGTAAACGTGGTT |
| Ccl2 | GCCTGCTGTTCACAGTTGC | CAGGTGAGTGGGGCGTTA |
| Ccl3 | AGATTCCACGCCAATTCATCG | GCCGGTTTCTCTTAGTCAGGA |
| Ccl4 | GCCCTCTCTCTCCTCTTGC | GAGAAACAGCAGGAAGTGGGA |
| Cd14 | AGCCTTTCTCGGAGCCTATC | AGCAACAAGCCAAGCACAC |
| Cd52 | GCCCAGGAAGATTTCAGGAT | CCCAAGGATCCTGTTTGTATCT |
| Cd68 | CTGTTCACCTTGACCTGCT | TCACGGTTGCAAGAGAAACA |
| Cd83 | TCCAGCTCCTGTTTCTAGGC | GGACACTGCATAGGAGAGCT |
| Cgas | TGGCAGCTACTATGAACATGTGA | CCTGGGGACTTCCAGTTTAAAC |
| Clec7a | AGAGTGAAGGGCCATGGTT | CCTGGGGAGCTGTATTTCTGA |
| Clu | AGCTGGCTAACCTCACAC | GACCTCTGAGTCAGAGGAATG |
| Csf1 | CCTAGTCTTGCTGACTGTTGG | CCAATGTCTGAGGGTCTCG |
| Ctsb | AAGCTGTGTGGCACTGTCC | GATCTATGTCCTCACCGAACGC |
| Cx3cr1 | GGCCTAGAGCTCAAAGAAATCC | CACAGACCTTCGATCCCAGT |
| Cxcl10 | GGGAAGCTTGAAATCATCCCT | TCAGACATCTCTGCTCATCATTC |
| Cxcl11 | TGCTGAGATGAACAGGAAGG | CGCCCCTGTTTGAACATAAG |
| Cxcl13 | TGAGGCTCAGCACAGCAA | ATGGGCTTCCAGAATACCGTG |
| Cxcl3 | GCCACACTCCAGCCTAGC | GCCACAACAGCCCCTGTA |
| Cxcl9 | CCATGAAGTCCGCTGTTCTT | GCATTCCTTATCACTAGGGTTCC |
| Cxcr3 | ACAAGTGCCAAAGGCAGAG | GGCATCTAGCACTTGACGTTC |
| Ddx58 | GCAGAACTGGAACAGGTCG | GTTCGAAGTCCGGGATGC |
| Egln3 | CCGGCTGGGCAAATACTATG | CCACATGGCGAACATAACCTG |
| Eif2ak2 | GTTGTTGGGAGGGAGTTGAC | AGAGGCACCGGGTTTTGTA |
| Emp1 | TTCGTGTTCCAGCTCTTCAC | CTGCTGGAGTTGAAGTTCCC |
| Fbln5 | AGGAAGATGGCATTCACTGCAG | GGCTGGTTCACACACTCGT |
| Gapdh | CAAGGTCATCCCAGAGCTGAA | CAGATCCACGACGGACACA |
| Gbp2 | GTAGACCAAAAGTTCCAGACAG | GATAAAGGCATCTCGCTTGG |
| Gfap | AGAACAACCTGGCTGCGTATA | CAGCGATTCAACCTTTCTCTCC |
| Ggta1 | AGATCGCATTGAAGAGCCTCA | ACGGGGTCACTGTCAAAACA |
| Grin1 | GATGGCAAGTTTGGCACACA | AGCAGCTCTCCCATCATTCC |
| Grin2a | AGGAGGAGTTTGTGGACCAA | CAAATCGGAAAGGCGGAGAA |
| Grin2b | TGCATCCGAAGCTGGTGATA | TGCAGGGACTTGTCTTTCCA |
| Grin2c | CGTAGACAGAGCAACCACAC | CGATGCAGAAGCCCTTACAA |
| Grn | GGGAAATCCTGCTTCCAGATG | TGGCAGAGTCAGGACATTCA |
| Gsk3b | CCCTCAAATCAAGGCACATCC | AGGTGTGTACTCCAGCAGAC |
| H2-D1 | AGGAACCTGCTCGGCTACTA | CCCAAGTCACAGCCAGACA |
| H2-T23 | ACGGCTGGGAAATGAGACA | GCACCTCAGGGTGACTTCA |
| Hmox1 | CCTCACAGATGGCGTCACTT | GCTGATCTGGGGTTTCCCTC |

| | | |
|---|---|---|
| Icam1 | TTGGAGCTAGCGGACCAG | GGACCGGAGCTGAAAAGTTG |
| Ifih1 | CTTGTCACGAACGAGATAGCC | CCAGGACATACGTGCTTTCAT |
| Ifnb1 | CGGACTTCAAGATCCCTATGG | ACCCAGTGCTGGAGAAATTG |
| Ifng | ACGGCACAGTCATTGAAAGC | TGTCACCATCCTTTTGCCAG |
| Il18 | ACAGCCTGTGTTCGAGGAT | TCACAGCCAGTCCTCTTACTT |
| Il1a | GGTTAAATGACCTGCAACAGGA | GAGCGCTCACGAACAGTT |
| Il1b | GCCACCTTTTGACAGTGATGAG | ACAGCCCAGGTCAAAGGTT |
| Il1r1 | TGGAAGGGATGACTATGTTGG | TGAAGCCTCCCATATCTCTCA |
| Il1rn | TGTGCCAAGTCTGGAGATG | GCGCTTGTCTTCTTCTTTGTTC |
| Il33 | GGTGAACATGAGTCCCATCA | CGTCACCCCTTTGAAGCT |
| IL6 | CGATGATGCACTTGCAGAAA | ACTCCAGAAGACCAGAGGAA |
| Irf7 | CTTCAGCACTTTCTTCCGAGA | TGTAGTGTGGTGACCCTTG |
| Irf8 | GAGCCAGATCCTCCCTGAC | GGCATATCCGGTCACCAGT |
| Itgam | GACTCTCATGCCTCCTTTGG | GTGGGTCCTGGACATGTTG |
| Itgax | TGGCTGTAGATGACCAAACGT | TGTGGTCAGCTCCACAGTT |
| Itgb2 | CATCCATGTGGAGGACAGTCT | CCAATCAGTACGACACCTACCA |
| Myd88 | GCCTTGTTAGACCGTGAGGA | CCTGGTTCTGCTGCTTACCT |
| Nlrc4 | GGCCTGCAACCTCTTTCTTA | CAGGTCTTCTTCTGTGACCTG |
| Nlrp12 | CACCAGACCTGCAGACTC | CATGCTTTGGAGGTGAGTCC |
| Nlrp1a | CCCGCTATATCGTGTCTTCC | CGGTAGCACAGCTCTAGTTC |
| Nlrp3 | TTCCCAGACACTCATGTTGC | AGAAGAGACCACGGCAGAA |
| Nos1 | GTCAAGTACGCCACCAACAA | GGAAGTCATGCTTGCCATCA |
| Nos2 | CTTTGCCACGGACGAGAC | TCATTGTACTCTGAGGGCTGAC |
| Pycard | GGAGCTCACAATGACTGTGCT | CTGCCACAGCTCCAGACTC |
| Rsad2 | CGAGGACTGCTTCTGCTCA | CCAAGTATTCACCCCTGTCCT |
| Serping1 | TGGCCCAATTCGATGACCATA | ACGGGTACCACGATCACAAA |
| Stat3 | CCCCGTACCTGAAGACCAA | ACACTCCGAGGTCAGATCCA |
| Sting1 | CCAACAGCGTCTACGAGATTC | ACATGGCAAACAGGGTCTG |
| Tbp | CCTTGTACCCTTCACCAATGAC | ACAGCCAAGATTCACGGTAGA |
| Timp1 | ATGCCCACAAGTCCCAGAAC | TGCAGGCACTGATGTGCAAA |
| Tlr2 | GGGCTTCACTTCTCTGCTT | AGCATCCTCTGAGATTTGACG |
| Tlr3 | GATACAGGGATTGCACCCATAA | TCCCCCAAAGGAGTACATTAGA |
| Tlr4 | GGACTCTGATCATGGCACTG | CTGATCCATGCATTGGTAGGT |
| Tlr9 | GGAGAATCCTCCATCTCCCAA | AGAGTCTCAGCCAGCACT |
| Tnf | GGGTGATCGGTCCCCAAA | TGAGGGTCTGGGCCATAGAA |
| Tnfrsf1b | GAAGGCTCAGATGTGCTGT | CCGAGGTCTTGTTGCAGAA |
| Trem2 | TGGGACCTCTCCACCAGTT | TGGTGTTGAGGGCTTGGG |
| Tyrobp | TGGTGTTGACTCTGCTGATTG | GTCTCAGCAATGTGTTGTTTCC |
| Vim | GATTTCTCTGCCTCTGCCAAC | CAACCAGAGGAAGTGACTCCA |
| VSV | TGATACAGTACA ATTATTTGGGAC | GAGACTTTCTGTTACGGGATCTGG |


1. Consensus recommendations for the postmortem diagnosis of Alzheimer's disease. The National Institute on Aging, and Reagan Institute Working Group on Diagnostic Criteria for the Neuropathological Assessment of Alzheimer's Disease. *Neurobiol Aging* **18**, S1-2 (1997).
